# Lamin A/C functions independently from mechanical signaling during adipogenesis

**DOI:** 10.1101/2020.09.07.279828

**Authors:** Matthew Goelzer, Amel Dudakovic, Melis Olcum, Buer Sen, Engin Ozcivici, Janet Rubin, Andre J van Wijnen, Gunes Uzer

**Affiliations:** Boise State University; Mayo Clinic; University of North Carolina Chapel Hill; Izmir Institute of Technology

**Keywords:** Lamin A/C, LINC, Nucleoskeleton, Nuclear Envelope, Adipogenesis, Mechanical Signals, Mesenchymal Stem Cells

## Abstract

Mesenchymal stem cells (MSC) maintain the musculoskeletal system by differentiating into multiple cell types including osteocytes and adipocytes. Mechanical signals, including strain and low intensity vibration (LIV), are important regulators of MSC differentiation. Lamin A/C is a vital protein for nuclear architecture that supports chromatin organization, as well as mechanical integrity and mechano-sensitivity of the nucleus in MSCs. Here, we investigated whether Lamin A/C and mechano-responsiveness are functionally coupled during adipogenesis. Lamin depletion in MSCs using siRNA increased nuclear area, height and volume and decreased circularity and stiffness, while phosphorylation of focal adhesions and dynamic substrate strain in response to LIV remained intact. Lamin A/C depletion decelerates adipogenesis as reflected by delayed appearance of key biomarkers (e.g., adiponectin/ADIPOQ). Based on RNA-seq data, reduced Lamin A/C levels decrease the activation of the adipocyte transcriptome that is normally observed in response to adipogenic cues mediating differentiation of MSCs. Mechanical stimulation via daily LIV application reduced the expression levels of ADIPOQ in both control and Lamin A/C depleted cells. Yet, treatment with LIV did not induce major transcriptome changes in either control or Lamin A/C depleted MSCs, suggesting that the biological effects of LIV on adipogenesis may not occur at the transcriptional level. We conclude that while Lamin A/C activation is essential for normal adipogenesis, it is dispensible for activation of focal adhesions by dynamic vibration induced mechanical signals.

## Introduction

As one of the Lamin family proteins that form the nucleoskeleton, Lamin A/C (gene symbol: LMNA) has a vital role in providing the mechanical and structural integrity of the cell nucleus [1–3]. Mutations in LMNA lead to premature aging in Hutchinson Gilford progeria syndrome [2–4], also known as progeria [1–3]. This mutation in LMNA causes alterations at histone methylation sites in heterochromatin [3, 4]. In pluripotent embryonic stem cells (ESCs), Lamin A/C protein is expressed at low basal levels in undifferentiated cells, but expression is elevated after differentiation into ESC derivatives [4–6]. Because Lamin A/C supports the formation of transcriptionally suppressed chromatin (i.e., heterochromatin), its low levels in ESCs is consistent with absence of heterochromatin in ESC cells [7]. In contrast, the nucleoskeleton protein Lamin B (encoded by the LMNB1 and LMNB2 genes) was found to be present both before and after differentiation. These studies collectively indicate that Lamin A/C plays a specific role during the differentiation of ESCs.

Mechanical and structural attributes of the cell and nucleus change during Lamin A/C loss [8]. When Lamin A/C is depleted, cellular elasticity and viscosity of the cytoplasm decreases [9]. Such a change in mechanical properties affects the response to external forces: the nucleus of Lamin A/C deficient cells display higher displacement magnitude than that of wild type cells in response to biaxial strain, indicating a lower nuclear stiffness [10]. In contrast, nuclei containing the progeroid farnesylated Lamin A/C (i.e. progerin) show increased stiffness when visualized under strain [11]. Both loss and mutation of Lamin A/C are associated with irregular nuclear morphology, including blebbing and loss of circularity [12, 13]. Additionally, because Lamin A/C is located at the inner nuclear membrane, it acts as an anchoring site for chromatin. Depletion of Lamin A/C has been shown to affect both the dynamics and the organization of the chromatin [14, 15], and may secondarily play a role in chromatin mediated mechanical properties of the cell nuclei [16]. Therefore, Lamin A/C plays a vital role in regulating cellular and nuclear mechanical structure and shape.

Mesenchymal Stem/Stromal Cells (MSCs) are tissue resident multipotent cells that can differentiate into musculoskeletal lineages including osteoblasts and adipocytes [17]. MSCs replace and rejuvenate skeletal and connective tissues in response to environmental mechanical demand, and their differentiation program is responsive to mechanical stimuli [18–21]. For example, application of external mechanical challenge in the form of LIV over 14-days increases proliferation and osteogenic differentiation markers and subsequent mineralization of MSC cultures *in vitro* [22–24]. In contrast to ESCs, MSCs are somatic cells that have the potential to differentiate into distinct mesenchymal lineages and express Lamin A/C in their native state. In this way, depletion of Lamin A/C in MSCs severely impedes osteoblast differentiation. MSCs treated with a siRNA targeting LMNA showed a drastic reduction in osteoblast differentiation transcription factors such as OCN, OSX, and BSP, and an increase in fat droplet formation when induced to differentiate into adipocytes [25]. Mutations of Lamin A/C, specifically the lipodystrophy-associated LMNA p.R482W mutation, can also serve to slow adipogenic differentiation in cells [26]. Additionally, overexpression of Lamin A/C has been shown to induce osteogenesis while inhibiting adipogenesis in human MSCs [27]. Mouse studies have shown that *Lmna* -/- mice have a significant reduction in bone mass compared to WT mice reflecting reduced osteoblast numbers [28]. While these findings suggest a role for Lamin A/C in regulating the differentiated state of MSCs, whether Lamin A/C depletion contributes to mechanical regulation of MSC differentiation remains insufficiently explored.

An important signaling node for mechanical control of MSC are focal adhesions, macromolecule protein complexes located on the cellular membrane, that connect the cytoskeleton to the extracellular matrix (ECM) where the cell is anchored to the extracellular environment through integrins [29]. During dynamic mechanical stimulus, integrin engagement is regulated by activation of Focal Adhesion Kinase (FAK) at tyrosine 397 residue [30]. We have reported both LIV and substrate strain lead to FAK phosphorylation at tyrosine 397[31]. This activation of FAK at focal adhesions both recruits signaling molecules that lead to cytoskeletal restructuring and activates concomitant mechanosignaling events such as the Akt/ß-Catenin (AKT1-CTNNB1) pathway [32]. Application of mechanical stimuli with strain, fluid flow, and LIV generates concomitant activation of ß-Catenin and RhoA signaling in MSCs [31, 33, 34]. Within the context of MSC adipogenesis, activation of these parallel signaling pathways results in decelerated adipogenic commitment of MSCs as measured by reduced production of adipogenesis related proteins such as adiponectin (encoded by adiponectin gene ADIPOQ) and peroxisome proliferator-activated receptor gamma (PPARG1) [35]. In addition to cytomechanical signaling events initiated at focal adhesions and cytoskeleton, control of MSC differentiation is also dependent on nuclear connectivity within the cytoskeleton. Inhibiting nucleo-cytoskeletal connectivity by disabling the function of Linker of Nucleoskeleton and Cytoskeleton (LINC) complexes impedes the nuclear entry of important molecular transducer mechanical information such as YAP/TAZ and ß-Catenin which act as co-transcriptional factors for regulating MSC adipogenesis and osteogenesis [36]. As opposed to LINC complex depletion, Lamin A/C depletion has no effect on mechanically induced nuclear ß-Catenin entry [37], suggesting that Lamin A/C may be dispensable for the mechanically-induced activation of focal adhesions that lead to de- phosphorylation and subsequent nuclear entry of ß-Catenin.

These previous studies show that Lamin A/C plays a central role in nuclear organization and structure, as well as contributing to the cell’s ability to sense structural qualities of the extracellular matrix to guide differentiation of MSCs. However, the role of Lamin A/C in focal adhesion signaling and mechanically-induced control of MSC fate in response to dynamic mechanical challenges remains incompletely understood. Therefore, we tested the requirement of Lamin A/C for the mechanical response of MSCs. Using LIV, we have investigated the role of Lamin A/C depletion on the mechanical control of MSC adipogenesis.

## Results

### siRNA depletion of Lamin A/C weakens the nuclear elastic modulus in MSCs

We investigated the effects of Lamin A/C loss on cellular and nuclear morphology as well as mechanical properties. MSCs treated with either a control siRNA (siCntl) or a Lamin A/C specific siRNA (siLMNA) were stained against F-actin and DNA. Compared to the siCntl group, siLMNA treated MSCs showed a more elongated nuclear morphology but no apparent changes in the F-actin cytoskeleton (**Fig. 1A**). Shown in **Fig. 1B**, morphology quantification indicated a 9% decrease of nuclear sphericity in siLMNA treated MSCs when compared to MSCs treated with a control siRNA (p<0.001). The nuclear area, volume, and height were increased 32%, 31%, and 11% in siLMNA treated MSCs, respectively, compared to control siRNA treated cells (**Fig. 1B**, p<0.001). The Young’s modulus was measured for both whole cells and extracted nuclei treated with either siLMNA or siCntl. The Young’s modulus was measured using a rounded AFM probe tip which was pressed onto the surface of the whole cell directly above the nucleus or on an isolated nucleus (**Fig. 1C** and **1D**). Confocal imaging with DNA and Lamin A/C labeling of a representative isolated nucleus (**Fig. 1E**), indicates that nuclear structure remains intact following isolation. Treatment with siLMNA caused a 45% reduction in whole cell stiffness when compared to siCntl treated MSCs (**Fig. 1F**, p<0.001), while extracted nuclei exhibit a 55% reduction in stiffness in Lamin A/C depleted cells compared to siCntl (p<0.01)(**Fig. 1G**).

**Fig. 1.**
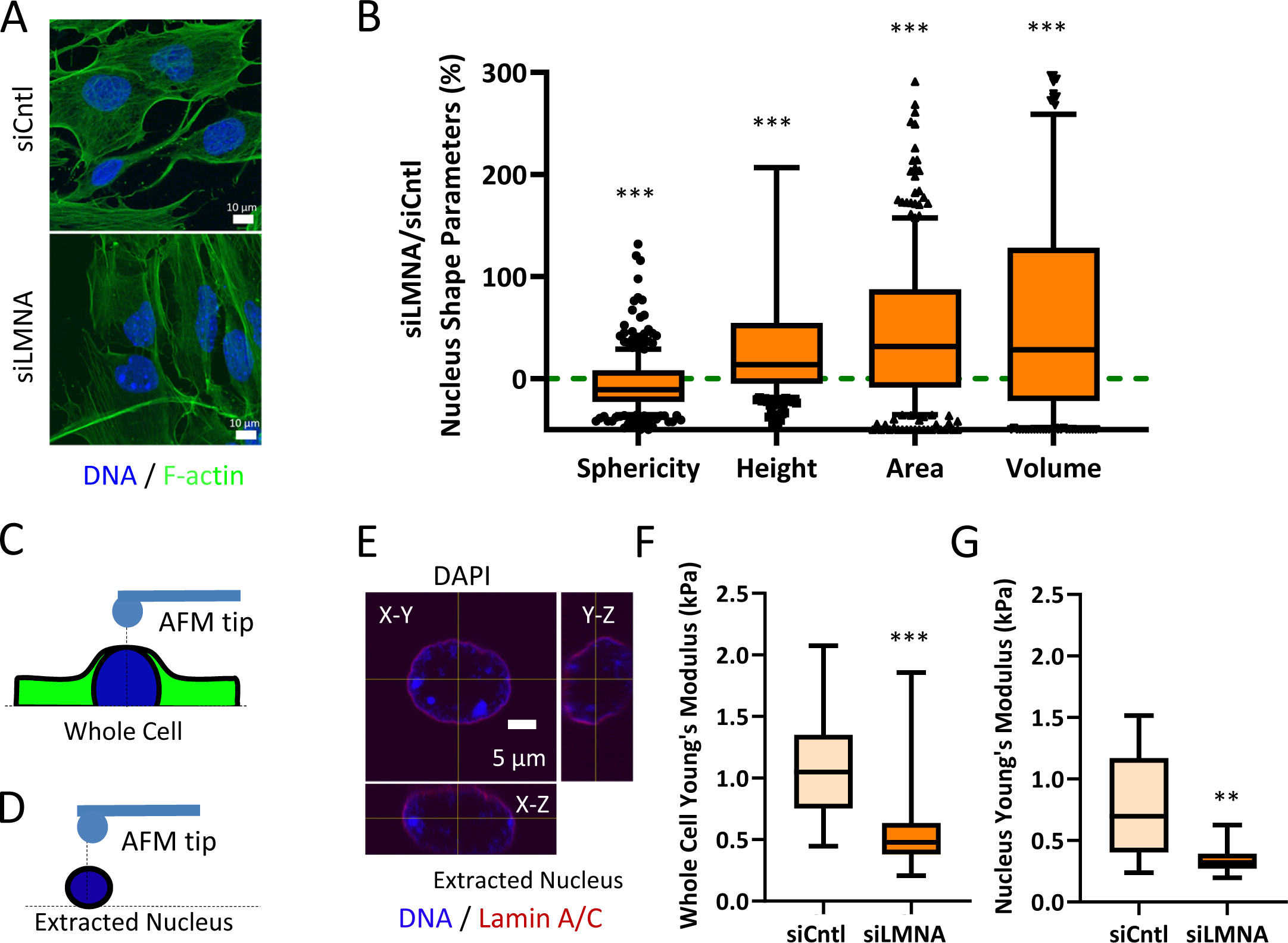
siRNA depletion of Lamin A/C weakens the nuclear elastic modulus in MSCs: (A) Confocal Image of F-actin (phallodin, green) and nucleus (DAPI, blue). Scale bar: 10µm. **(B)** Geometric parameters of siCntl and siLMNA groups quantified and presented as a % difference compared to siCntl group (green line). Nuclear sphericity decreased by 8% in MSCs treated with Lamin A/C specific siRNA (siLMNA) compared to MSCs treated with a non-specific control siRNA (siCntl) (p<0.05, n=342). Nuclear area of siLMNA treated cells showed a 32% increase when compared to siCntl (p<0.05, n=342). Nuclear volume siLMNA treated cells increased by 31% compared to siCntl (p<0.05, n=342). Nuclear height of siCntl and siLMNA treated cells. When compared to the nuclear height of siCntl MSCs, siLMNA treated cells had increased nuclear height of 12% (p<0.05, n=342). **(C)** Schematic of AFM probe tip testing whole cell Young’s modulus in live MSCs. **(D)** Depiction of AFM probe tip testing live extracted nucleus. **(E)** Confocal image of extracted nucleus depicting its orthogonal views from X-Y, X-Z, Y-Z planes (DAPI, blue; Lamin A/C, Red) Scale bar: 5µm. **(F)** Whole cell Young’s modulus of the siLMNA group was 45% lower when compared to the siCntl group. **(G)** Young’s modulus of extracted live nucleus in siLMNA MSCs remained 55% lower when compared to siCntl MSCs (p<0.01, n=13). Results are presented as mean ± STD. Group comparisons were made via non- parametric Mann Whitney U-test. p<0.05, ** p<0.01, *** p<0.001, against control.

### siRNA depletion of Lamin A/C (LMNA) Increases Sun-2 (SUN2) Nuclear Levels and Focal Adhesion Proteins

To further characterize the effects of Lamin A/C loss on nuclear envelope and focal adhesions, the LINC complex and focal adhesion proteins were investigated. Confocal images of the siCntl and siLMNA groups indicated that there were no visible changes in the LINC proteins Sun-1 (SUN1) and Sun-2 (SUN2) when Lamin A/C was depleted (**Fig. 2A**). Quantitative analysis of the confocal images did not detect any differences in Sun-1 or Sun-2 nuclear envelope localization (**Fig. S1**). We examined the same proteins using cellular fractionation followed by western blotting and densitometry analysis (**Fig 2B)**. All the measurements were normalized to whole cell siCntl protein amounts which was set to 1. Comparing siLMNA treatment with siCntl, Lamin A/C significantly decreased in whole cell Lamin A/C (-35%, p < 0.05). The relative Lamin A/C concentration was greater in the nuclear fraction and led to larger values, while band intensities of the siLMNA group remained significantly lower compared to siCntl (-11%, p < 0.05). Except for a small amount of Sun-1 detection in the cytoplasm, both Sun-1 and Sun-2 were largely restricted to the nucleus. Knocking down Lamin A/C was associated with an increase in nuclear Sun-2 (+44%, p<0.05). Focal adhesion proteins also were altered under siLMNA treatment. Total focal adhesion kinase (FAK) adhered to the cell culture plate experienced an increase of 39% compared to control treated cells (p<0.05) (Shown in **Fig. 2D** and **Fig. 2E**). The amount of Akt adhering to cell culture plates also increased by 50% (p<0.05). No changes in vinculin were detected.

**Fig. 2.**
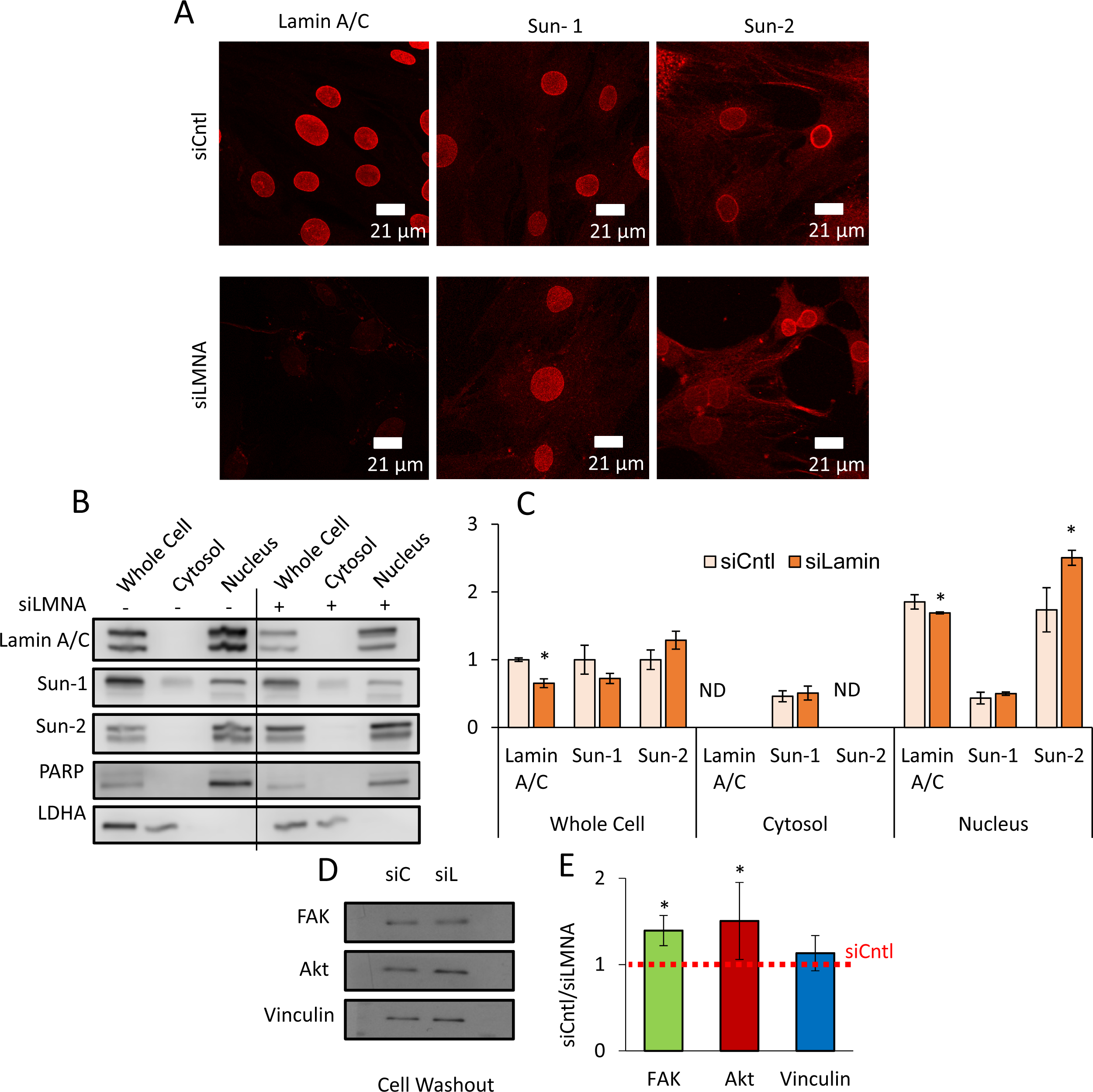
siRNA depletion of Lamin A/C affects the Sun-1 and Sun-2 elements of the LINC Complex and Focal Adhesion Proteins: (A) Confocal images of cells treated with the siCntl and siLMNA siRNA groups. Primary antibodies targeted Lamin A/C, Sun-1, and Sun-2. **(B)** Representative western blots of cell fractionations (whole cell, cytosol and nucleus) with cells treated with either siCntl or siLMNA. Primary antibodies targeted Lamin A/C, Sun-1, Sun-2, PARP, and LDHA. Line represents removal of protein ladder marker lane, uncropped blots are provided in Fig.S1. **(C)** Analysis of western of cell fractionation western blots (n=3/grp). siLMNA treated cells had 29% increase of Sun-2 in whole cells, 122% in cytoplasm, and 44% increase in nucleus fraction (p<0.05) compared to siCntl samples. Sun-1 levels saw a decrease of 28% in whole cell, 10% increase in cytoplasm, and 15% increase in nucleus fraction compared to siCntl samples. ND represents non-detectable levels. **(D)** Representative western blot of focal adhesion proteins following a cell washout. Primary antibodies targeted of FAK, Akt, and Vinculin in siCntl and siLMNA siRNA treated cells. **(E)** Densitometry analysis showed that, when compared to siCntl levels siLMNA treated MSCs showed increased levels of total FAK (39%, p<0.05) and total Akt (50%, p<0.05), no change in Vinculin was detected (n=3/grp). Results are presented as mean ± STE. Scale bar: 21µm. Group comparisons were made via parametric two-tailed Student T-test (C) or one-way ANOVA followed by a Newman-Keuls post-hoc test (E). p<0.05, ** p<0.01, *** p<0.001, against control.

### Focal adhesions maintain response to mechanical stimulus in Lamin A/C depleted MSCs

Basal levels of FAK were increased in Lamin A/C depleted cells. We next asked if mechanical activation of FAK was altered by further quantifying the mechanical activation of FAK via its phosphorylation at Tyrosine 397 residue (pFAK) which is indicative of integrin engagement [30]. MSCs were treated with either strain or LIV and compared to non-mechanically stimulated controls. Basal pFAK levels normalized to total FAK (TFAK) were 85% elevated in the siLMNA groups when compared to the siCtnl groups (p<0.05) (**Fig.3A** and **Fig.3B**). Phosphorylated FAK levels from both siCtnl and siLMNA treated groups increased by 101% (p<0.05) and 87% (p<0.001) in response to 20 min strain (2%, 0.1Hz) when compared to non-strained counterparts. LIV also activated FAK: pFAK increased by 331% (p<0.001) in siCtnl and 83% (p<0.001) in siLMNA treated MSCs in response to LIV (0.7g, 90Hz).

**Fig. 3.**
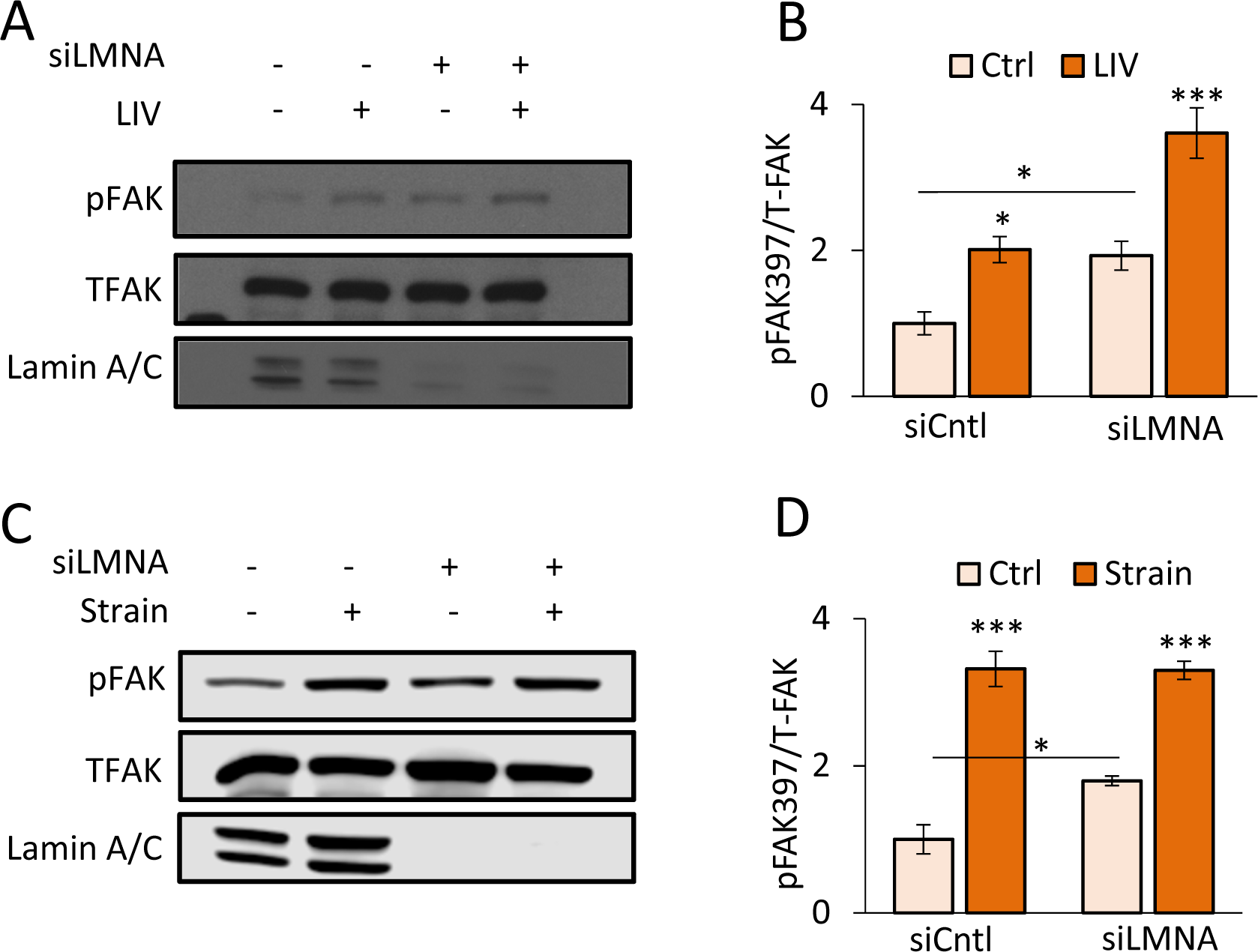
**Focal adhesions maintain response to mechanical stimulus in Lamin A/C depleted MSCs**: **(A)** Representative western blots for pFAK (Tyr 397), TFAK, and Lamin A/C in siCntl and siLMNA treated cells groups treated with 2 bouts of LIV (20min, 90Hz, 0.7g) separated by 2 hour rest period. LIV treated sample had a 2-fold increase of pFAK compared to non-LIV. **(B)** Analysis of western image of pFAK, TFAK, and Lamin A/C during LIV (n=4/grp). The non-LIV siLMNA group had a 92% (p<0.05) increased basal pFAK compared to the non-LIV siCntl group. In response to LIV, both siCtnl and siLMNA treated MSCs elicited 101% (p<0.05) and 87% increases in pFAK, respectively. **(C)** Representative western blots for pFAK (Tyr 397), TFAK, and Lamin A/C of the siCntl and siLMNA groups treated with a single bout strain (20 min, 0.1 Hz, 2% strain). **(D)** Analysis of pFAK, TFAK, and Lamin A/C immediately after strain application (n=4/grp). The non-strain siLMNA group had a 79% (p<0.05) increased basal pFAK presented as mean ± STE. Group comparisons were made via one-way ANOVA followed by a Newman-Keuls post-hoc test. p<0.05, ** p<0.01, *** p<0.001, against control or against each other.

### Application of daily LIV treatment decreases adipogenic differentiation in MSCs

As focal adhesion signaling was intact in siLMNA treated MSCs, we next probed downstream processes to ask whether the LIV application known to slow adipogenesis [38] was effective when Lamin A/C was depleted. In our experiment timeline, cells were first treated with siRNA on day 1 and then cultured in adipogenic media concomitant with LIV treatment (**Fig. 4A**). On day 2, adipogenic media was placed on cells and LIV treatment started. LIV treatment occurred twice a day for 20 minutes with two hour rests in between treatments. On day 7, cell protein or RNA samples were collected for either western blotting or RNA-seq analysis. Probing adipogenesis marker adiponectin between non-LIV controls, Lamin A/C depleted cells showed a 39% decrease in adiponectin protein at 7 days (**Fig. 4B** and **Fig. 4C)**. Similarly, compared within LIV treated groups, adiponectin levels in the siLMNA group was 51% lower than siCntl treated cells with LIV (p<0.01). Compared to non-LIV controls, daily LIV application decreased adiponectin protein levels by 30% in the siCntl (p<0.01) and 44% in the siLMNA groups (p<0.001).

**Fig. 4.**
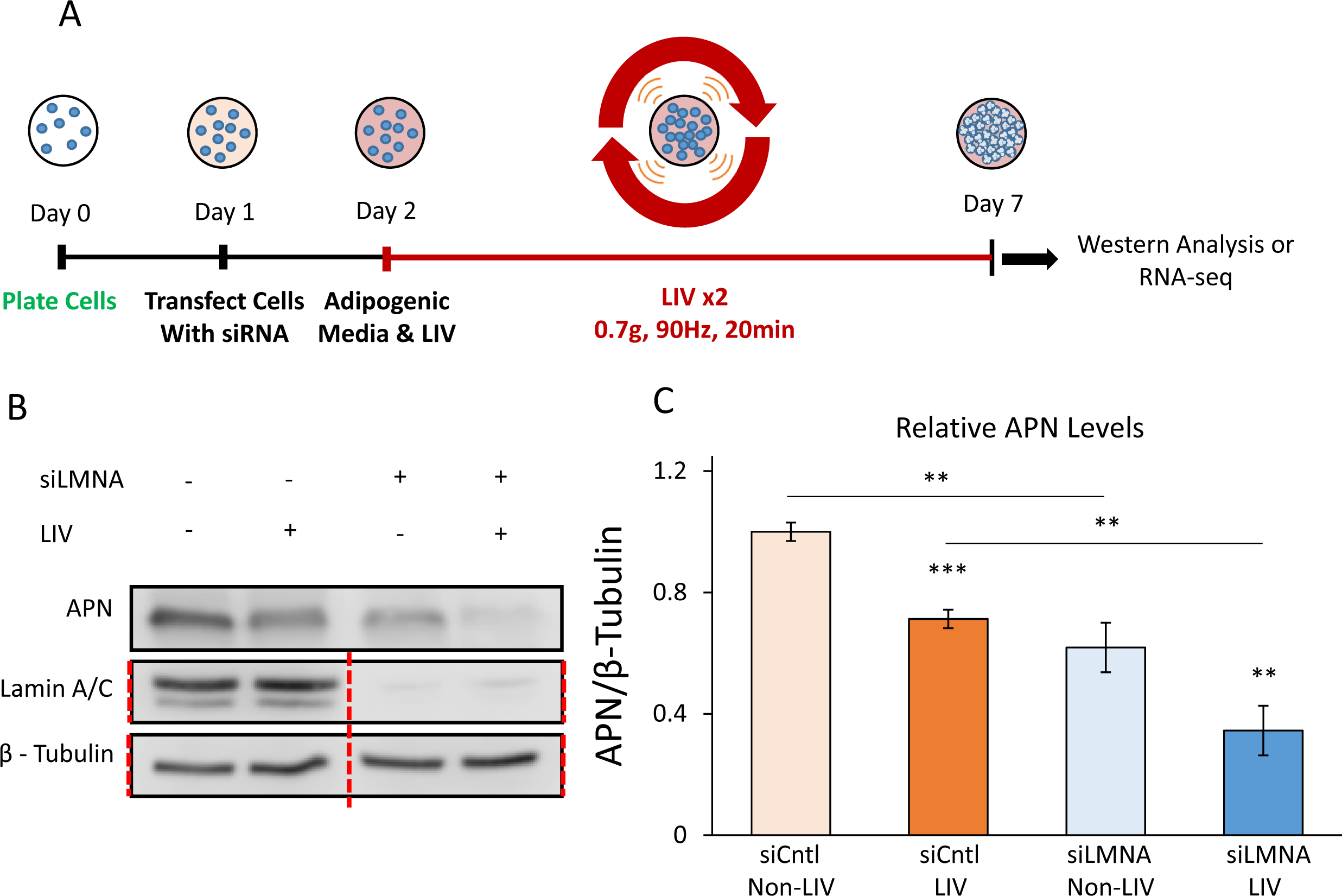
Application of daily LIV treatment decreases adipogenic differentiation in MSCs: (A) Timeline of experiments. On day 0 cells were plated on 10cm dishes. Then, on day 1 cells were transfected with siRNA. On day 2 adipogenic media was placed on cells and cells were treated with LIV for 20 minutes, twice daily. Once cells differentiated cells were pulled off for either western analysis or RNA-seq. **(B)** Representative western blots of cells treated with siCntl and siLMNA after 7 days of adipogenic induction with and without LIV treatment. Adiponectin protein, Lamin A/C, and β-Tubulin were targeted. Lamin A/C and β – Tubulin were imaged on the same plot. Red line represents western blot cropped for alignment; uncropped blots were provided in Fig.S4. **(C)** Relative levels of adiponectin of the siCntl and the siLMNA groups. Compared to siCntl MSCs with no LIV, adiponectin protein levels in siLMNA treated MSCs with no LIV were decreased by 39% (p<0.01, n=4). Compared to non-LIV controls for siCntl treated cells, LIV treated samples had 30% reduction in adiponectin protein levels (p < 0.001, n=3/grp). SiLMNA treated cells treated with LIV had a 44% reduction of Adiponectin protein compared to non-LIV samples (p<0.01, n=3/grp). Compared to siCntl cells with LIV treatment, siLMNA cells treated with LIV had a 51% reduction in adiponectin (p<0.01, n=3/grp). Results are presented as mean ± STE. Group comparisons were made via one-way ANOVA followed by a Newman-Keuls post-hoc test. p<0.05, ** p<0.01, *** p<0.001, against control or against each other.

### Differential effect of Lamin A/C depletion and LIV on mRNA transcription during adipogenic differentiation

RNA-seq was performed to determine the effects of LIV and siLMNA treatment on differential mRNA in MSCs during adipogenesis. Read values were filtered for robust expression by selecting genes with average levels of 0.3 FPKM (<u>Fr</u>agments <u>pe</u>r <u>k</u>ilobase of transcript per <u>m</u>illion mapped reads), t-test p < 0.05, and Log2 fold change greater than 1.4). Hierarchical clustering of these genes generated a heatmap (**Fig. 5A**) in which siCntl treated samples clustered together in one clade, while undifferentiated and siLMNA treated samples were clustered together in another clade that is visually separated from siCntl treated samples. Principal component analysis (**Fig. 5B**) shows further grouping of siCntl samples and siLMNA samples. Corresponding LMNA FPKM values were presented in **Fig. S2**. Principal component 1 and component 2 explain 40.4% and 15.9% total variance, respectively with prediction ellipses indicating the probability of 0.95 that a new observation of the same group will fall inside the ellipse. Representative RNA-seq data for individual genes shows FPKM levels for a panel of 13 genes associated with the adipogenic pathway, including adiponectin (ADIPOQ), CCAAT/enhancer-binding protein alpha (CEBPA), and peroxisome proliferator-activated receptor gamma (PPARG) and others (**Fig. 5C**).

**Fig. 5.**
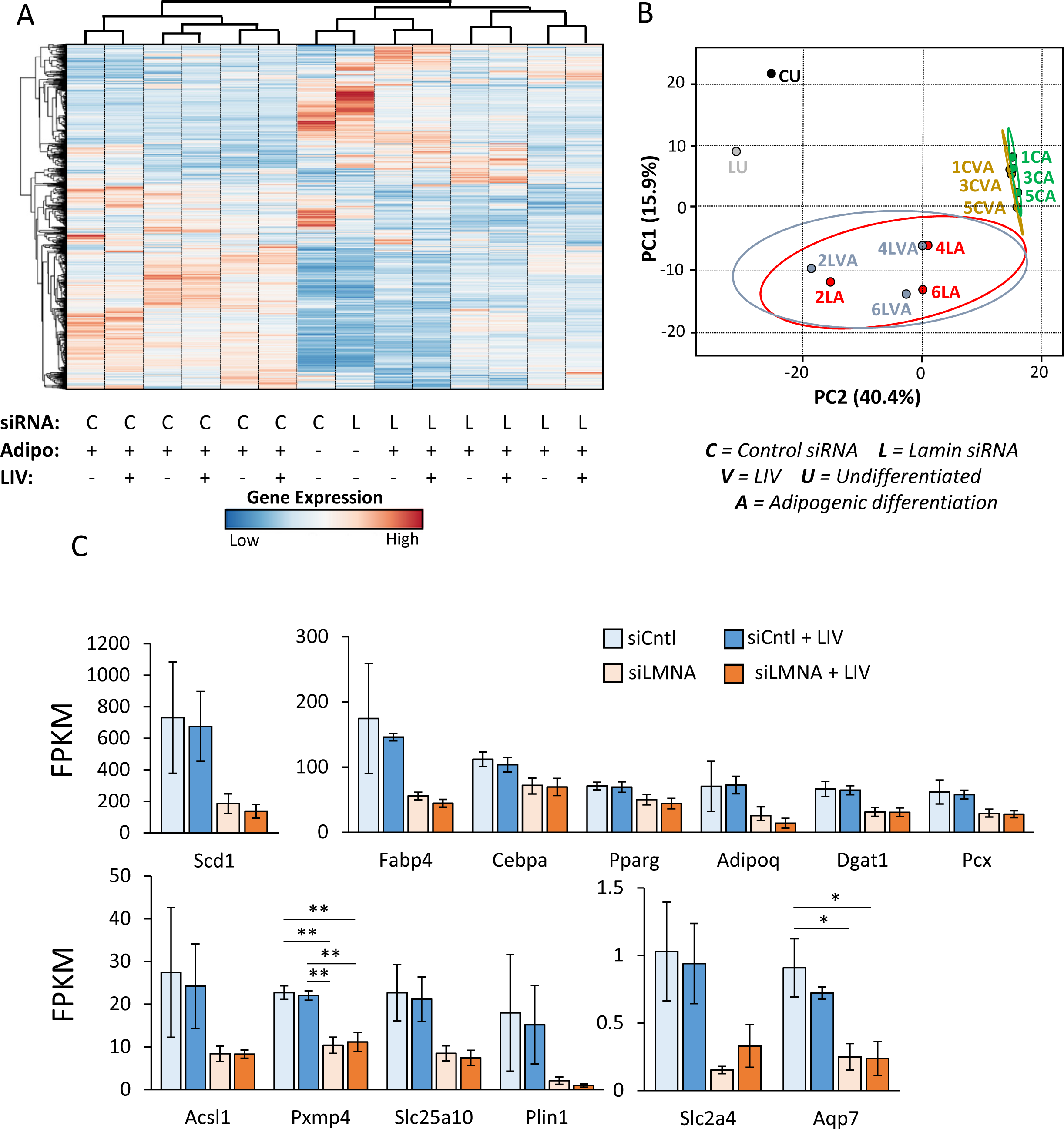
Differential effect of Lamin A/C depletion and LIV on MSCs transcription duringadipogenic differentiation: (A) Heat map of genes with average expression of 0.3 FPKM, t-test p < 0.05, and fold change greater than 1.4. Unit variance scaling is applied to rows. **(B)** Principle component plot where principal component 1 and principal component 2 that explain 40.4% and 15.9% of the total variance, respectively. Prediction ellipses are such that with probability 0.95, a new observation from the same group will fall inside the ellipse. N = 14 data points. **(C)** Average FPKM values of genes related to adipogenic phenotype. Results are presented as mean ± STE. Group comparisons were made via one-way ANOVA. * p<0.05, ** p<0.01 were against control or against each other.

### Lamin A/C depletion impedes adipogenic transcription in MSCs

Cells treated with siLMNA and siCntl with adipogenesis were compared statistically to determine differential gene expression between siRNA treatments. A volcano plot for the comparison between siLMNA and siCntl treated samples under adipogenic constraints (**Fig. 6A**) revealed there are 52,607 statistically unchanged transcripts between Lamin A/C depleted and control MSCs with Wald values of p>0.05 (grey and green data points). Shown in green data points, 2,000 of them showed at least a 2-fold difference (i.e. Log2 fold change ≥ 1). While 749 genes showed statistically significant change between Lamin A/C depleted and control MSCs with Wald values of p<0.05 (shown in blue) and 427 of them had a less than 2-fold difference (i.e. Log2 fold change ≤1). The remaining 322 genes showed at least a 2-fold difference (i.e. Log2 fold change ≥1), which represents significant and differentially expressed genes. Up-regulated (red genes on the right side, n = 173) and down-regulated genes (red genes on the left side, n = 149) upon LMNA depletion were then assessed by a clustering analysis using ClustVis [39]. Upregulated genes upon Lamin A/C depletion are associated with cellular processes such as (i) Tissue Repair (e.g., genes generally involved in angiogenesis, hematopoiesis, and mechanical stress shielding), (ii) ECM remodeling (e.g. genes generally involved in take-up and intra-cellular transport of ECM debris as well as suppression of apoptosis), (iii) cell surface transporters (e.g., genes that mediate the trafficking of compounds across membranes) (**Fig.6B**). Collectively, the biological function of these genes appear to be related to tissue repair, inflammation and extracellular matrix homeostasis. Downregulated gene groups upon LMNA knock-down included (i) Cell adhesion and cytoskeletal organization, (ii) interferon signaling and regulation of gene expression (e.g., DNA and RNA binding, and protein degradation), (iii) G protein coupled receptor signaling (e.g., diverse range of cell surface receptors and components of the angiotension system), (iv) lipid metabolism and paracrine inflammatory signaling, and (v) adipogenic phenotype (**Fig.6C**). These down regulated genes together are generally involved in cell migration, energy metabolism and adipogenic differentiation. The results from gene ontology and gene network analysis revealed that Lamin A/C depletion has pleiotropic effects on gene expression, yet many gene pathways converge on cell surface related biochemical events, interactions with the extracellular matrix and internal metabolic pathways.

**Fig. 6.**
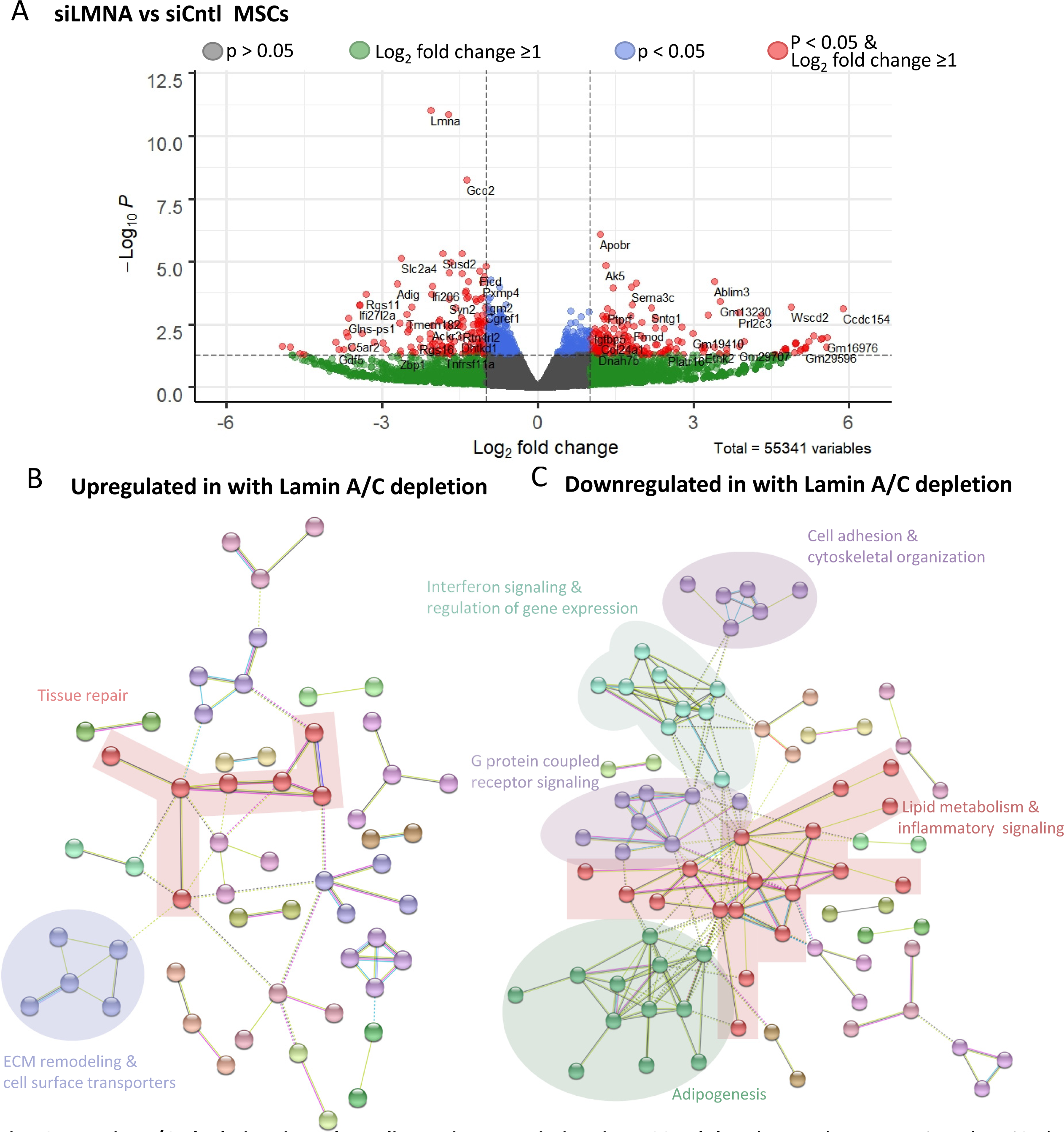
Lamin A/C depletion impedes adipogenic transcription in MSCs: (A) Volcano plot of siLMNA compared to siCntl under adipogenic conditions. Genes with Wald values of p>0.05 are colored in grey. Genes with 2-fold differential gene expression but have Wald values of p>0.05 are colored in green. Genes colored with blue have Wald values of p<0.05, but less than 2-fold differential gene expression. Genes with Wald values of p<0.05 and greater than 2-fold gene expression are colored in red. Grouping of five or more associated genes were highlighted and subsequently subjected to a supervised analysis of biologic function. **(B)** Upregulated genes were associated with cellular processes included tissue repair, ECM remodeling and cell surface transporters. Full size image is presented in Fig. S3 **(C)** Downregulated gene groups included, cell adhesion and cytoskeletal organization, interferon signaling and regulation of gene expression, G-protein coupled receptor signaling, lipid metabolism and paracrine inflammatory signaling and adipogenic phenotype. Full size image is presented in Fig. S4

### LIV Decreases Interferon Signaling Pathway in siLMNA and siCntl Treated Cells

To determine the effects of LIV with siCntl and siLMNA controls under adipogenic constraints were compared against their LIV treated counterparts. The volcano plot comparing the siCntl adipogenesis with or without LIV treatment (siCntl ± LIV) is shown in **Fig. 7A.** There were 53,326 statistically unchanged genes between with Wald values of p>0.05 (grey and green data points) with, 1,939 of them showed at least 2-fold difference (green). While 76 genes showed statistically significant change between LIV treated and control MSCs with Wald values of p<0.05, shown in blue, 26 of them had a less than 2-fold difference. Remaining 53 genes showed at least 2-fold difference (red). Assessing down-regulated genes via clustering revealed an interferon-related cluster in the LIV treatment group (**Fig. 7B**). Similarly, LIV treatment upon LMNA depletion also revealed an interferon-related cluster when assessing gene clustering by ClustVis in the significant and highly down-regulated genes (**Fig. 7D**). Together, cells treated with siCntl had 11 genes that are part of interferon pathway while cells treated with siLMNA had 16 genes associated with interferon pathway signaling (**Fig. 7E**). While interferons have been reported to have effects on adipogenesis of mouse embryonic fibroblasts [40], the physiological relevance of the interferon pathway is uncertain, because this pathway may be linked to cellular responses to events precipitated by siRNA transfection as well. Excluding this latter finding, it appears that while LMNA loss has a dramatic impact on gene expression programs, LIV has very minimal effects on the transcriptome of differentiating MSCs.

**Fig. 7.**
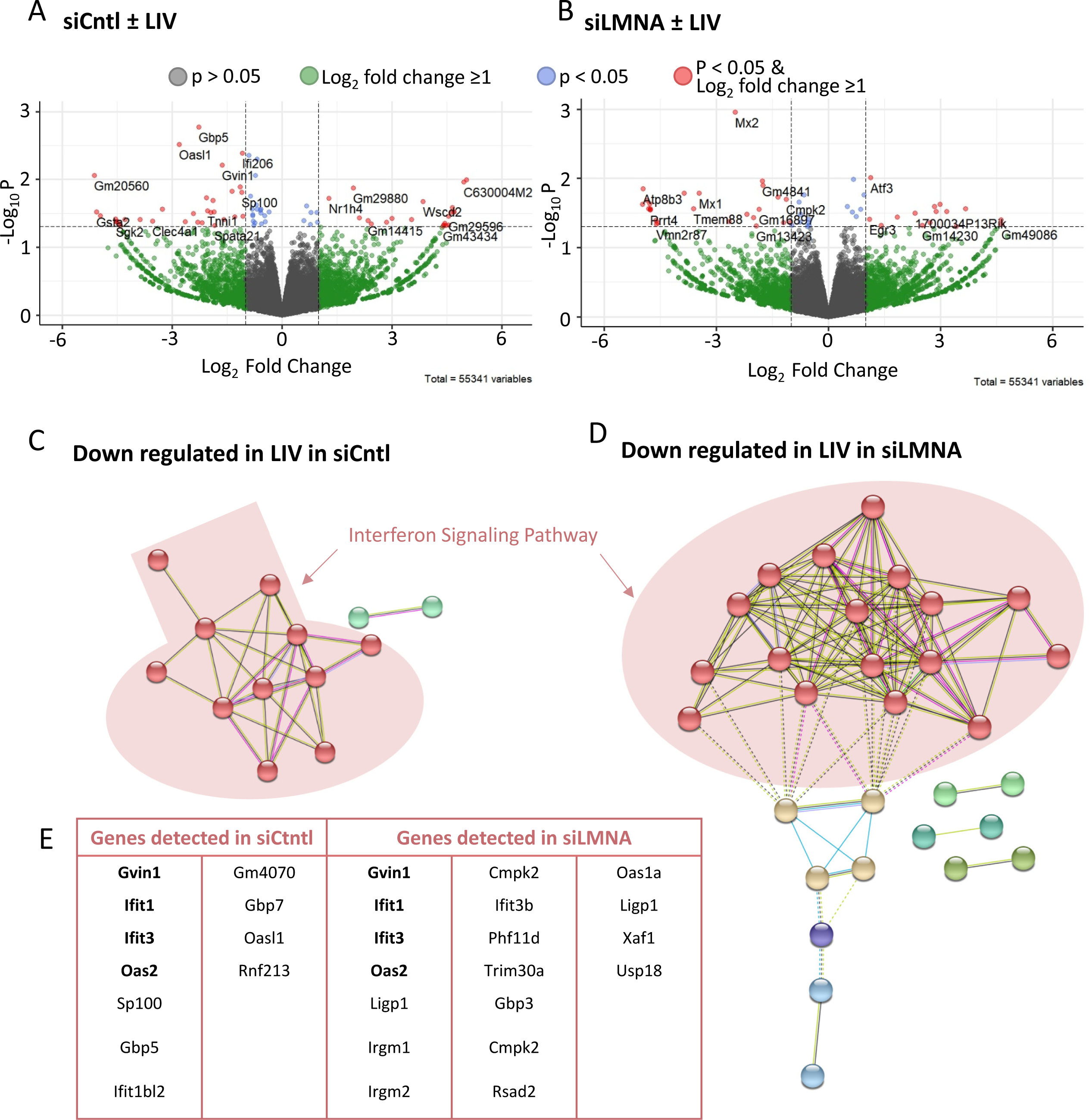
LIV Decreases Interferon Signaling Pathway in siLMNA and siCntl Treated Cells: (A) Volcano plot comparing the siCntl adipogenesis with or without LIV treatment (siCntl ± LIV). **(B)** Volcano plot comparing the siLMNA adipogenesis with or without LIV treatment (siLMNA ± expression but have Wald values of p>0.05 are colored in green. Genes colored with blue have Wald values of p<0.05, but less than 2-fold differential gene expression. Genes with Wald values of p<0.05 and greater than 2-fold gene expression are colored in red. Both siCtrl **(C)** and siLMNA **(D)** showed downregulation of genes closely associated with interferon signaling pathway. Full size images are presented in Fig. S5A and S5B. **(E)** Cells treated with siCntl had 11 genes associated with interferon signaling pathway while siLMNA treated cells had 16 genes associated with interferon pathway. **Bolded** gene names (Gvin, Ifit1, Ifit3, and Oas2) names were found in both siCntl and siLMNA treated samples.

## Materials and Methods

### MSC Isolation

Bone marrow derived MSC (mdMSC) from 8-10 wk male C57BL/6 mice were isolated as described [41]. Briefly, tibial and femoral marrow were collected in RPMI-1640, 9% FBS, 9% HS, 100 μg/ml pen/strep and 12μM L-glutamine. After 24 hours, non-adherent cells were removed by washing with phosphate-buffered saline and adherent cells cultured for 4 weeks. Passage 1 cells were collected after incubation with 0.25% trypsin/1 mM EDTA × 2 minutes, and re-plated in a single 175-cm2 flask. After 1-2 weeks, passage 2 cells were re-plated at 50 cells/cm^2^ in expansion medium (Iscove modified Dulbecco’s, 9% FBS, 9% HS, antibiotics, L-glutamine). mdMSC were re-plated every 1-2 weeks for two consecutive passages up to passage 5 and tested for osteogenic and adipogenic potential, and subsequently frozen.

### Cell Culture, Pharmacological Reagents, and Antibodies

Fetal calf serum (FCS) was obtained from Atlanta Biologicals (Atlanta, GA). Culture media, trypsin-EDTA, antibiotics, and Phalloidin-Alexa-488 were from Invitrogen (Carlsbad, CA). MSCs were maintained in IMDM with FBS (10%, v/v) and penicillin/streptomycin (100μg/ml). For phosphorylation measurements, seeding cell density was 10,000 cells per square centimeter. For immunostaining experiments, seeding cell density was 3,000 cells per square centimeter. For phosphorylation measurements and immunostaining experiments, all groups were cultured for 48h before beginning experiments and were serum starved overnight in serum free medium.

For adipogenic differentiation experiments, the seeding cell density was 21,000 cells per square centimeter. Cells were transfected 24 hours after cell seeding with siRNA targeting Lamin A/C (siLMNA) or a control sequence (siCntl) using RNAiMax from Invitrogen. Adipogenic media and LIV treatment followed previously published protocol, where twenty four hours after the transfection, the adipogenic media was added which contained dexamethasone (0.1μM) and insulin (5 μg/ml) [18]. Cell cultures were incubated with the combined transfection media and adipogenic differentiation media for 7 days after adipogenic media was added with or without LIV treatment (2 X 20 minutes per day separated by 2 hours).

The following antibodies were purchased: Cell Signaling (Danvers, MA): Akt (#4685), p-Akt Ser473 (#4058L), β-Tubulin (D3U1W), and p-FAK Tyr397 (#328 3). ThermoFischer Scientific (Rockford, Il): Adiponectin (PA1-054). Santa Cruz Biotechnology (Dallas, TX): FAK (sc-558), Lamin A/C (sc-7292).

### LIV and Strain

Vibrations were applied at peak magnitudes of 0.7g at 90Hz twice for 20min separated by 2h rest period at room temperature. Uniform 2% biaxial strain was delivered at 10 cycles per minute for 20 min using the Flexcell FX-5000 system (Flexcell International, Hillsborough, NC). Controls were sham handled. During adipogenesis experiments, LIV was applied 24 hours after initial transfection, a regimen we previously shown be effective [31].

### Isolation of Focal Adhesions

Cells were incubated with triethanolamine (TEA)-containing low ionic-strength buffer (2.5 mM TEA, pH 7.0) for 3 minutes at RT, 1× PBS containing protease/phosphatase inhibitors. A Waterpik (Fort Collins, CO, www.waterpik.com) nozzle held 0.5 cm from the plate surface at approximately 90° supplied the hydrodynamic force to flush away cell bodies, membrane-bound organelles, nuclei, cytoskeleton, and soluble cytoplasmic materials so that residual focal adhesions could be isolated as we have reported previously[31] .

### siRNA Silencing Sequences

For transient silencing of MSCs, cells were transfected with gene-specific small interfering RNA (siRNA) or control siRNA (20 nM) using RNAiMax (ThermoFischer) according to manufacturer’s instructions. The following Stealth Select siRNAs (Invitrogen) were used in this study: LaminA/C 5′-UGGGAGAGGCUAAGAAGCAGCUUCA-3′ and negative control for LaminA/C 5′- UGGGAGUCGGAAGAAGACUCGAUCA-3′.

### Isolation of Nuclei for Young’s modulus

MSCs were plated at 10,000 cell/cm^2^ cell density. For mechanical and structural testing, nuclei were isolated by scraping cells in PBS and then suspending cells in hypotonic solution followed by centrifugation at 3000xg. Nuclei were then extracted by using percol (81% percol, 19% hypotonic buffer) and centrifugation at 10,000xg. Nuclei were then diluted in PBS and plated. Nuclei Young’s modulus was determined using Atomic Force Microscopy (AFM). For strain experiments, cells were plated on Bioflex Collagen-I coated silicone plates.

### RNA-seq

RNA extraction and sequencing were done by Novogene. Quality control of raw data was done using FASTQC. Read Alignment of the genome to the raw reads was done using STAR [42]. Read count generation was generated using feature Counts and Differential gene expression analysis was done using DESEQ2 [43]. For analysis using fragments per kilobase of transcript per million mapped reads (FPKM), data were assessed as previoulsy described [44–47]. Briefly, RNA-Seq data were analyzed by a Mayo Bioinformatics Core called MAPRSeq v.1.2.1 [48], which includes TopHat 2.0.6 alignment [49] with and gene expression quantification using HTSeq software [50]. Normalized gene counts were obtained from MAPRSeq as FPKM. Hierchical clustering and principal component analysis were assessed and visualized using ClustVis [39]. RNA-Seq data were deposited in the Gene Expression Omnibus of the National Institute for Biotechnology Information (GSE157056).

### Immunofluorescence

Twenty four hours after the siRNA treatment against Lamin A/C protein, cells were fixed with 4% paraformaldehyde. Cells were permeabilized by incubation with 0.3% Triton X-100. Cells were incubated in a blocking serum in PBS with 5% Donkey Serum (017-000-121, Jackson Immuno Research Laboratories). Primary antibody solution were incubated on the cells for 1h at 37°C, followed by secondary antibody incubation of either Alexa Flour 594 goat anti-rabbit (Invitrogen) or Alexa Fluor 647 donkey anti-mouse. For nuclear staining cells were incubated with NucBlue Hoeschst stain (Fischer Scientific). For actin staining, cells were incubated in Alexa Fluor 488 Phalloidin (Life Technologies). Primary and secondary concentrations were both 1:300.

### Nuclear Morphology

To test the nuclear morphology that will show the level of mechanical constraint on nucleus, MSCs seeded at 3000cell/cm^2^ on plastic slide chambers (iBIDI µslide # 80421). 72h after the siRNA treatment against Lamin A/C protein, DNA (Hoechst 33342, Life Technologies), and or immunostained against actin (Alexa Fluor 488 Phalloidin, Life Technologies). Z-stack confocal 3D images were obtained with a Zeiss LSM 710 with a separation interval of 0.15µm. Z-stack images were analyzed using IMARIS software.

### Western Blotting

Whole cell lysates were prepared using an radio immunoprecipitation assay (RIPA) lysis buffer (150mM NaCl, 50mM Tris HCl, 1mM EDTA, 0.24% sodium deoxycholate,1% Igepal, pH 7.5) to protect the samples from protein degradation NaF (25mM), Na3VO4 (2mM), aprotinin, leupeptin, pepstatin, and phenylmethylsulfonylfluoride (PMSF) were added to the lysis buffer. Whole cell lysates (15μg) were separated on 10% polyacrylamide gels and transferred to polyvinylidene difluoride (PVDF) membranes. Membranes were blocked with milk (5%, w/v) diluted in Tris-buffered saline containing Tween20 (TBS-T, 0.05%). Blots were then incubated overnight at 4°C with appropriate primary antibodies. Following primary antibody incubation, blots were washed and incubated with horseradish peroxidase-conjugated secondary antibody diluted at 1: 5,000 (Cell Signaling) at RT for 1h in 5% milk in TBST-T. Chemiluminescence was detected with ECL plus (Amersham Biosciences, Piscataway, NJ). At least three separate experiments were used for densitometry analyses of western blots and densitometry was performed via NIH ImageJ software.

### Statistical analysis

Results for densitometry were presented as mean ± SEM. Densitometry and other analyses were performed on at least three separate experiments. Differences between groups were identified by two-tailed Student’s T-test. Analysis of nuclear morphology and Young’s modulus were done using Whitney-Mann test and results were presented as mean ± STD. Differential gene expression analysis was done using Wald test. P-values of less than 0.05 were considered significant.

## Discussion

In this study, we have found that Lamin A/C depleted MSCs were able to activate focal adhesion signaling and decrease the output of adipogenic biomarkers (e.g., adiponectin) as efficiently as MSCs with intact Lamin A/C in response to LIV. Our findings indicate that the global adipogenic mRNA repression in Lamin A/C depleted MSCs occurred independent of LIV. RNA-seq analysis showed that LIV had negligible effects on mRNA levels compared to Lamin A/C depletion, suggesting that LIV effects on adipogenesis is likely caused by post-translational mechanisms or other downstream effects.

Lamin A/C depletion interfered with adipogenic differentiation but not with biomechanical responses. Not only was Lamin A/C dispensable for the LIV and strain mediated activation of focal adhesions, but LIV decreased levels of adiponectin protein. Consistent with repression of adipogenesis, in a Lamin A/C independent fashion. Adipogenic mRNA levels determined by RNA-seq were unaffected by LIV suggesting that LIV-induced repression of adipogenesis was post-transcriptional or post-translational.

In Lamin A/C depleted cells, microscopic observations of increased blebbing, elongated nuclear shape, and ruffled nuclear membrane [8, 12, 51] indicates a compromised nuclear structure. Quantification of 3D nuclear structure of Lamin A/C depleted cells were supportive of these previous observations and showed reduced sphericity and increased planar nuclear area while nuclear height and volume were increased compared to controls. It has been reported that Lamin A/C depletion increases nuclear height and volume in-part due to reduced recruitment of perinuclear and apical F-actin cables [52]. While reduction of apical F-actin may contribute to a decrease in elastic modulus in Lamin A/C depleted intact MSCs, a similar decrease was observed in Lamin A/C depleted isolated live nuclei. The similarities in decreased stiffness in both intact cells and isolated nuclei suggests that nuclear softening is the primary driver of decreased cell stiffness upon Lamin A/C loss of function.

Our data suggest that MSCs compensate for Lamin A/C mediated nuclear softening by increasing their focal adhesions. Not only was total FAK (PTK2) and Akt (AKT1) accumulation at the focal adhesions more robust in Lamin A/C depleted MSCs, Tyrosine 397 phosphorylated FAK was also higher which suggests increased integrin engagement [30]. These findings are not surprising as both depletion of Lamin A/C [53] and nucleo-cytoskeletal connector Nesprin-1 [54] were shown to increase substrate traction in cells. Tracking with increased basal pFAK levels, application of either LIV or strain pushed acute FAK phosphorylation of Lamin A/C depleted cells higher than control cells. These results indicate that the focal adhesion signaling remains intact in Lamin A/C depleted MSCs.

Similar to focal adhesions, nucleo-cytoskeletal connectivity provided by LINC complex remained intact under Lamin A/C depletion. Previous studies have shown that LINC proteins Sun-1 and Sun-2 bind to the Lamin A/C in order to mediate a connection from the inner nucleus to the cytoskeleton and ultimately to the focal adhesions that make a physical connection to the extra cellular matrix [55, 56]. Quantification of confocal images of Sun-1 and Sun-2 revealed no changes compared to controls while Sun-2 had an increase in proteins levels in both the whole cell and nuclear fractions. These observed protein changes under loss of the Lamin A/C could be in parallel to increased focal adhesion presence. Therefore, the cell may be increasing the levels of Sun-2 that is connected to actin, which in turn are connected to a higher number of focal adhesions. Localization of Sun-1 and Sun-2 proteins to the nuclear envelope are not entirely dependent upon Lamin A/C, but loss of Lamin A/C still results in some alteration of Sun-1 localization and no alteration for Sun-2 supporting previous literature as seen in **Fig. 2C** in the whole cell and nuclear fractions [56, 57] . While noted changes in the Sun proteins under Lamin A/C depletion suggests a putative relationship, loss of Lamin A/C did not negatively impact the structural Sun-mediated integrity of the LINC complex.

Adipogenesis has recently been shown to decrease with mutated Lamin A/C, specifically in cells expressing the lipodystrophy-associated LMNA p.R482W mutation [26]. Our data supports this previous observation as MSCs treated with siLMNA experienced slower adipogenic differentiation compared to siCntl treated cells (**Fig. 4C**). This observation is in contrast to studies that showed increased adipogenesis in Lamin depleted MSCs [25, 27, 58]. While cell culture conditions vary from experiment to experiment, this study did not utilize strong adipogenic inducers such as indomethacin and IBMX [27, 58]. Instead, we used a milder adipogenic media incorporating insulin and dexamethasone. This selection was based on previous work where LIV was unable overcome the adipogenesis induced by indomethacin and IBMX [18]. RNA-seq data indicate that upon MSCs display an undifferentiated phenotype upon Lamin A/C depletion, as reflected by reduced expression of genes associated with adipogenic and lipid related metabolic pathways. In contrast, LIV treatment did not have a significant impact on adipogenic gene expression indicating that Lamin A/C and not low intensity vibration is critical for adipogenic differentiation.

In contrast to large shifts in transcription under Lamin A/C depletion, RNA-seq data indicates that only 21 genes for siCntl and 74 genes for siLMNA treated cells were differentially expressed as a result of LIV treatment. Despite the lack of changes at mRNA level, Lamin A/C depleted MSCs retain their ability to respond to mechanical signals and exhibit decelerated adipogenesis reflected by reduced adiponectin protein in LIV treated cells. Although mechanical stimulation using LIV is not causing widespread alteration in mRNA expression, we did observe a distinct LIV-dependent signature characteristic of interferon responsive genes. Changes in interferon responses could be expected, because siRNA transfection affects single and double-stranded RNA accumulation within cells that may provoke interferon responses by mimicking viral RNA transfection. As such, this finding could perhaps be dismissed as technical artefact. However, this interferon related differentially expressed gene changes were seen as compared to non-LIV siRNAs. Hence, a biological cause may also be entertained. A possible relationship between type 1 interferon signaling pathway and the known mechanosensitive Wnt/β-catenin signaling pathway has been proposed [59]. GSK-3β is known to activate type 1 interferon signaling pathway [60] and inhibit the Wnt/β-catenin pathway by causing the degradation of β-catenin [37]. Activation of the Wnt/β- catenin signaling pathway via mechanical stimulus causes GSK-3β to be inhibited, promoting β- catenin translocation to the nucleus to inhibit adipogenesis [37], and potentially inhibit the type 1 interferon signaling pathway. Additionally, mechanical forces, specifically low intensity forces such as shear strain and vibration, have been shown to inactivate interferons [61]. Thus, there may be secondary mechanisms by which interferons respond to mechanical forces. The more important finding is the absence of major transcriptome changes during adipogenesis in response to LIV which points to post-transcriptional or post-translational regulatory events. While the mechanism of the observed mechanoregulation of adipogenesis is beyond of scope of this paper, further research will be needed to fully understand the potential mechanoregulation of adipogenesis during or after transcription.

## Conclusion

Lamin A/C depletion resulted in decreased nuclear integrity, more robust focal adhesions, and reduced adiponectin protein levels. Neither Sun-mediated LINC connectivity nor focal adhesion signaling in response to acute mechanical challenge were negatively impacted by Lamin A/C depletion. This independence of mechanical signaling from Lamin A/C was further highlighted by the significant reduction in adiponectin protein levels in response to LIV. The small transcriptional response under LIV was dwarfed by large transcriptional changes and blunted adipogenesis under Lamin A/C depletion. Findings of this study indicate that Lamin A/C is required for proper adipogenic commitment of MSCs into the adipogenic lineage and that the mechanical regulation of adipogenesis may not utilize similar pathways to elicit a response in MSCs.

## Data availability

The datasets generated and/or analyzed during the current study are available from the corresponding author on reasonable request.

## Acknowledgements

This study was supported by NIH AG059923 (GU), NSF 1929188 (GU), P20GM109095 (GU), R01AR049069 (AJvW), AR066616 (JR), Career Development Award in Orthopedics Research (AD), The Scientific and Technological Research Council of Turkey 2214-A (MO).

## Contributions

Matthew Goelzer: concept/design, data analysis/interpretation, manuscript writing

Amel Dudakovic: manuscript writing, data analysis, final approval of manuscript

Melis Olcum: concept/design, data analysis/interpretation, final approval of manuscript

Buer Sen: data analysis, final approval of manuscript

Engin Ozcivici: final approval of manuscript

Janet Rubin: data analysis, final approval of manuscriptAndre van Wijnen: manuscript writing, data analysis, final approval of manuscript

Gunes Uzer: concept/design, data analysis/interpretation, financial support, manuscript writing, final approval of manuscript

---

**Fig. S1.**
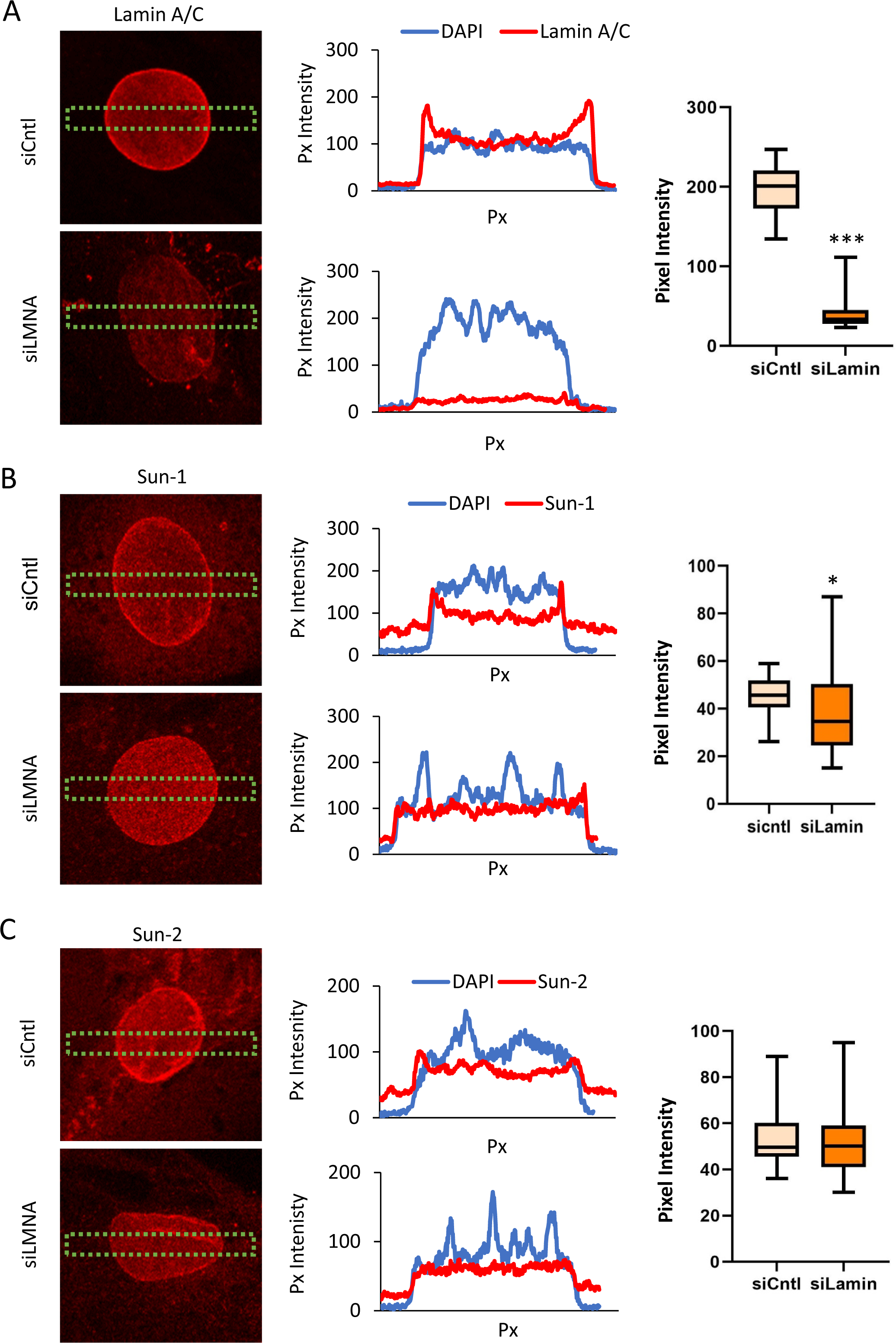
(A) Intensity profile of Lamin A/C staining along a rectangular region of interest on the nucleus. The middle plot shows the representative intensity distribution of Lamin A/C over the nucleus (blue, Hoechst 33342). Lamin A/C intensity peaked at the nuclear rim in siCntl cells while no peaks were observed in siLMNA cells. Comparison of peak intensity values at the nuclear envelope show 80% (p<0.001, n=25/grp) decrease with siLMNA treatment. **(B)** Intensity profile of Sun-1 staining along a rectangular region of interest on the nucleus. The middle plot shows the representative intensity distribution of Sun-1 (red) over the nucleus (blue, Hoechst 33342). Comparison of peak intensity values at the nuclear envelope show 15% (p<0.05, n=19/grp) decrease with siLMNA treatment. **(C)** Intensity profile of Sun-2 staining along a rectangular region of interest on the nucleus. The middle plot shows the representative intensity distribution of Sun-2 (red) over the nucleus (blue, Hoechst 33342). No difference between siCntl and siLMNA was detected. Images were quantified using ImageJ. Results are presented as mean ± SD. Group comparisons were made via non-parametric Mann Whitney U- test. * p<0.05, ** p<0.01, *** p<0.001.

**Fig. S2.**
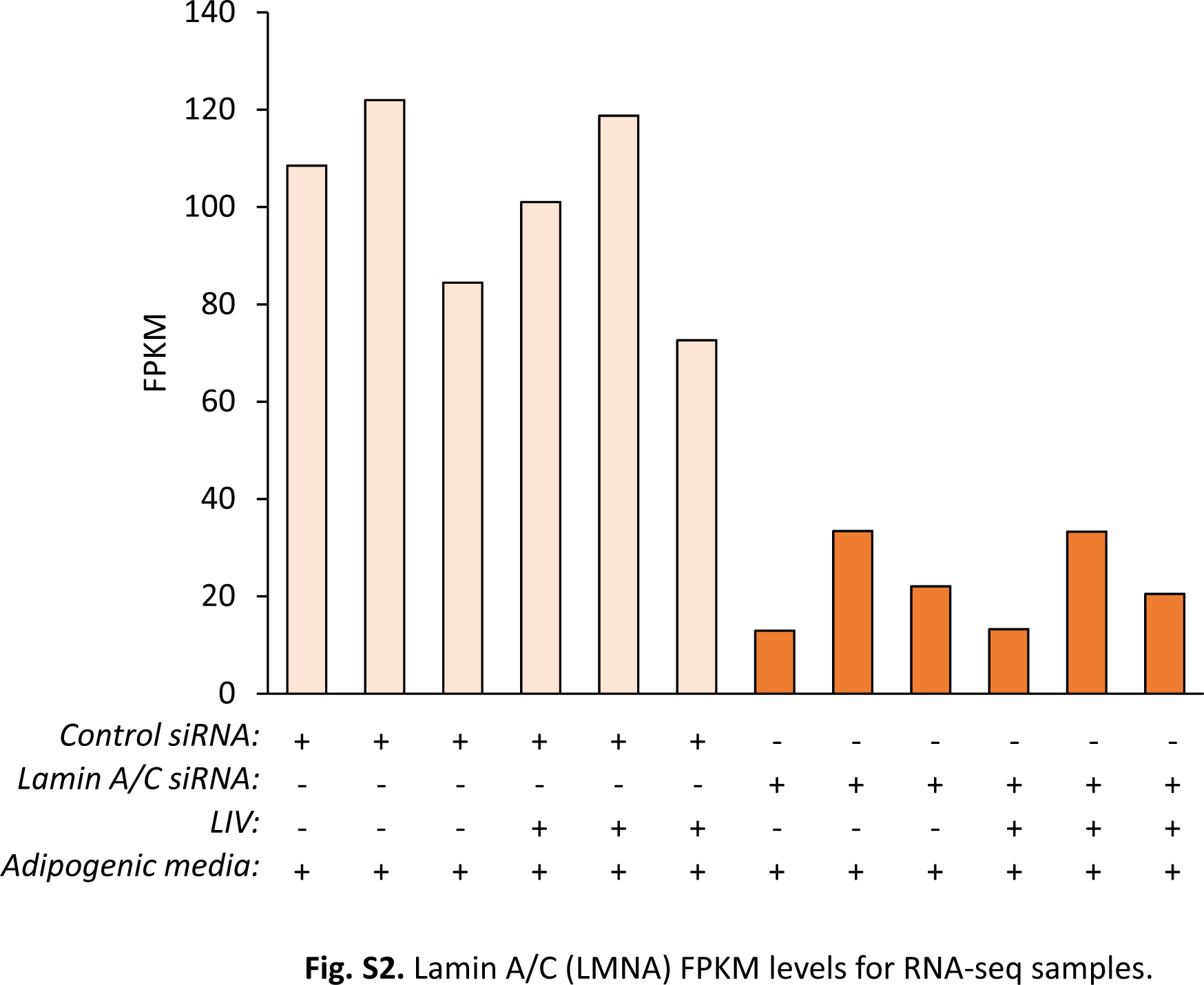
Lamin A/C (LMNA) FPKM levels for RNA-seq samples.

**Fig. S3.**
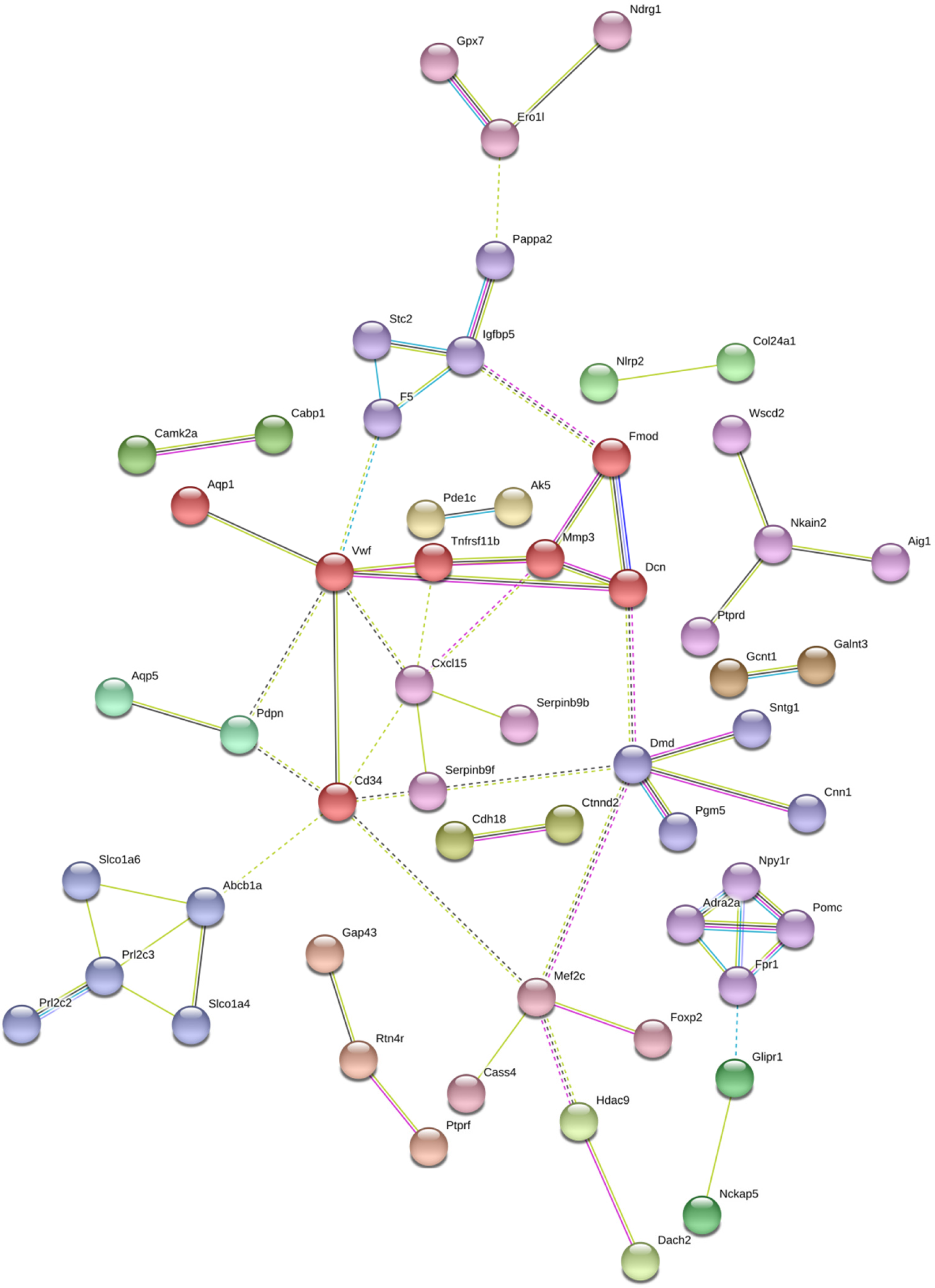
Full size, annotated gene cluster for Fig.6B

**Fig. S4.**
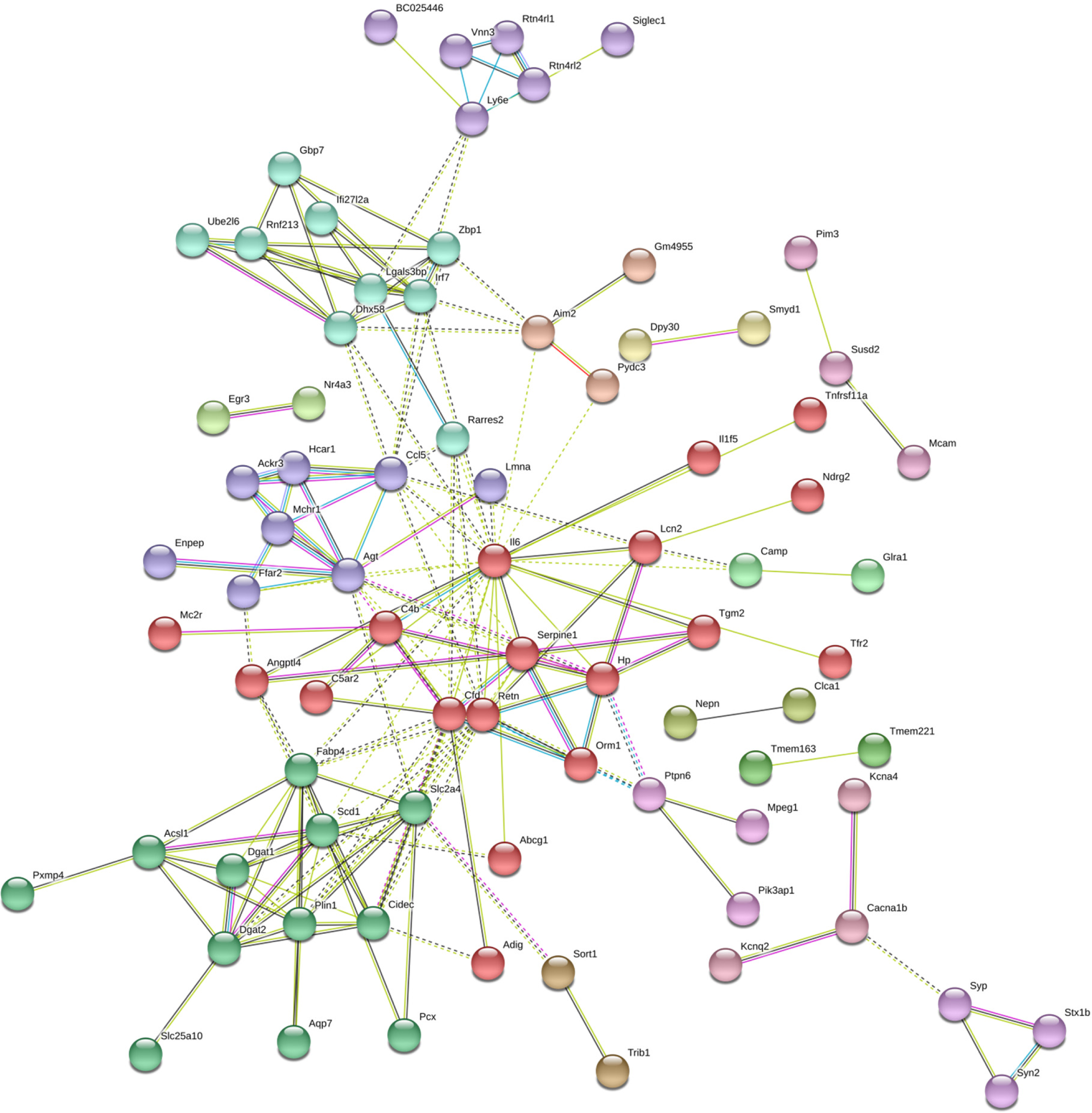
Full size, annotated gene cluster for Fig.6D

**Fig. S5.**
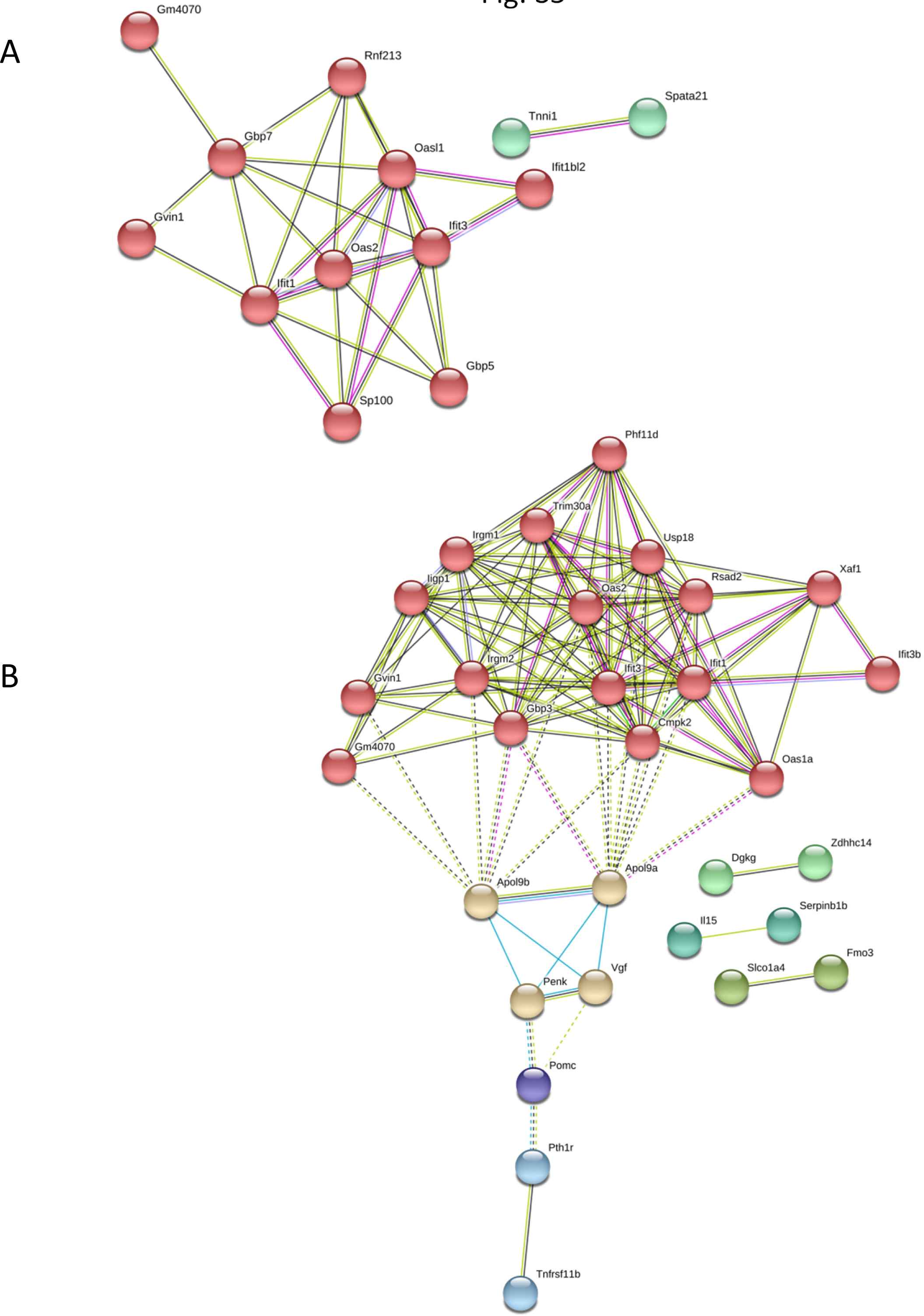
Full size, annotated gene cluster for (A) Fig.7C and (B) Fig.7D

**Fig. S6.**
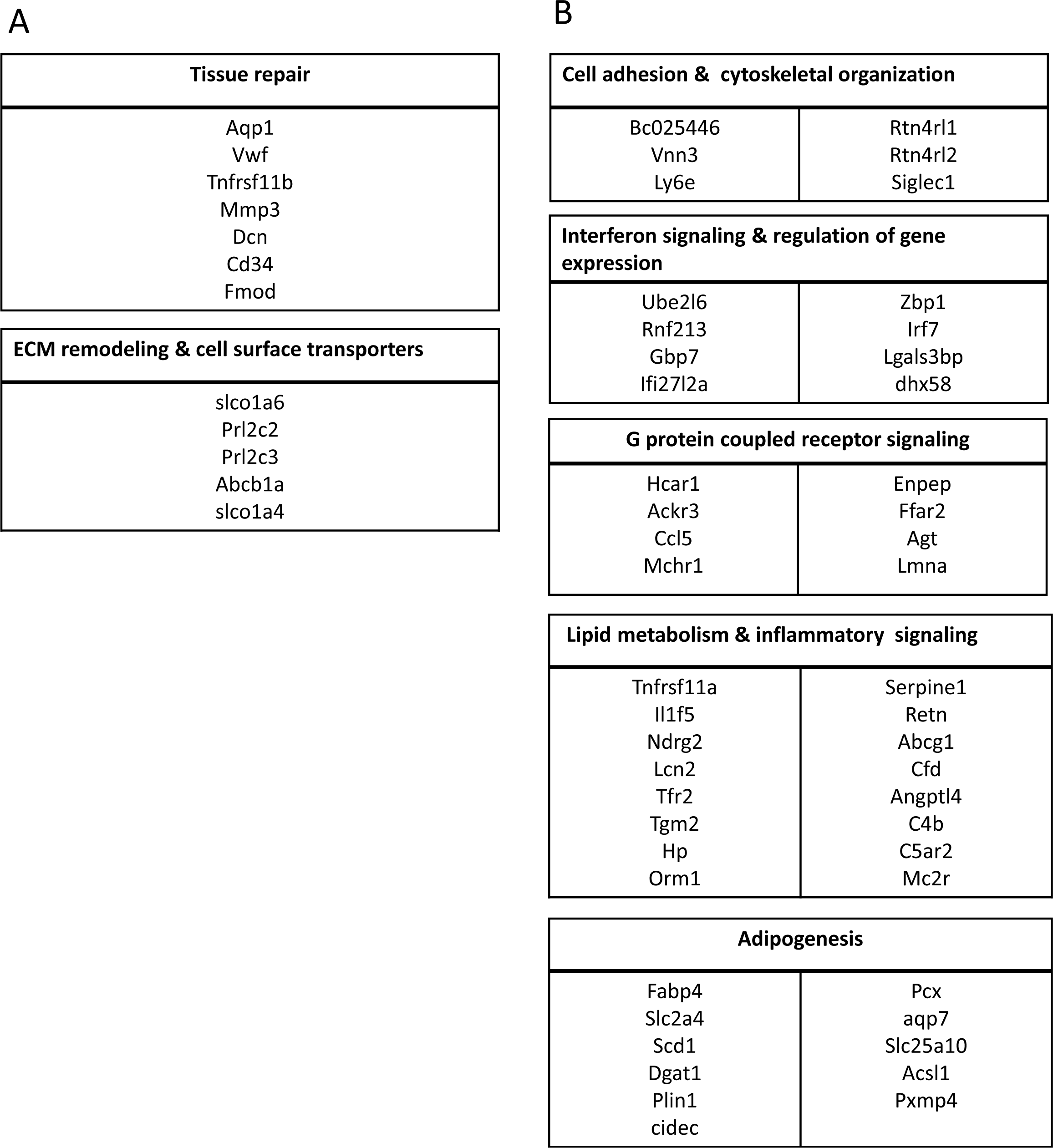
Gene lists for (A) Fig.6B and (B) Fig.6C

**Fig. S7.**
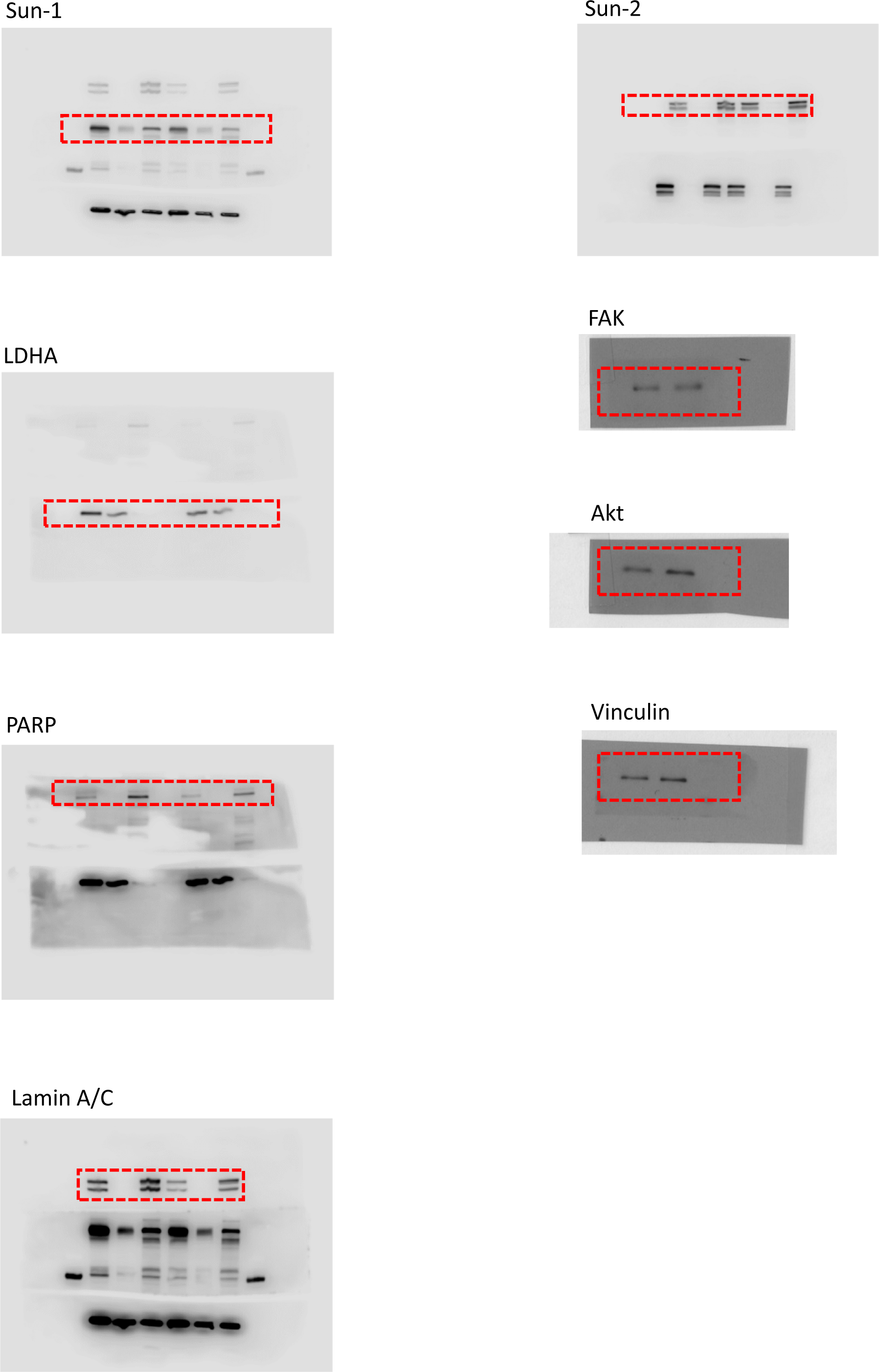
Unprocessed blots used in Figure 2 as obtained by LiCor C-DiGit blot scanner.

**Fig. S8.**
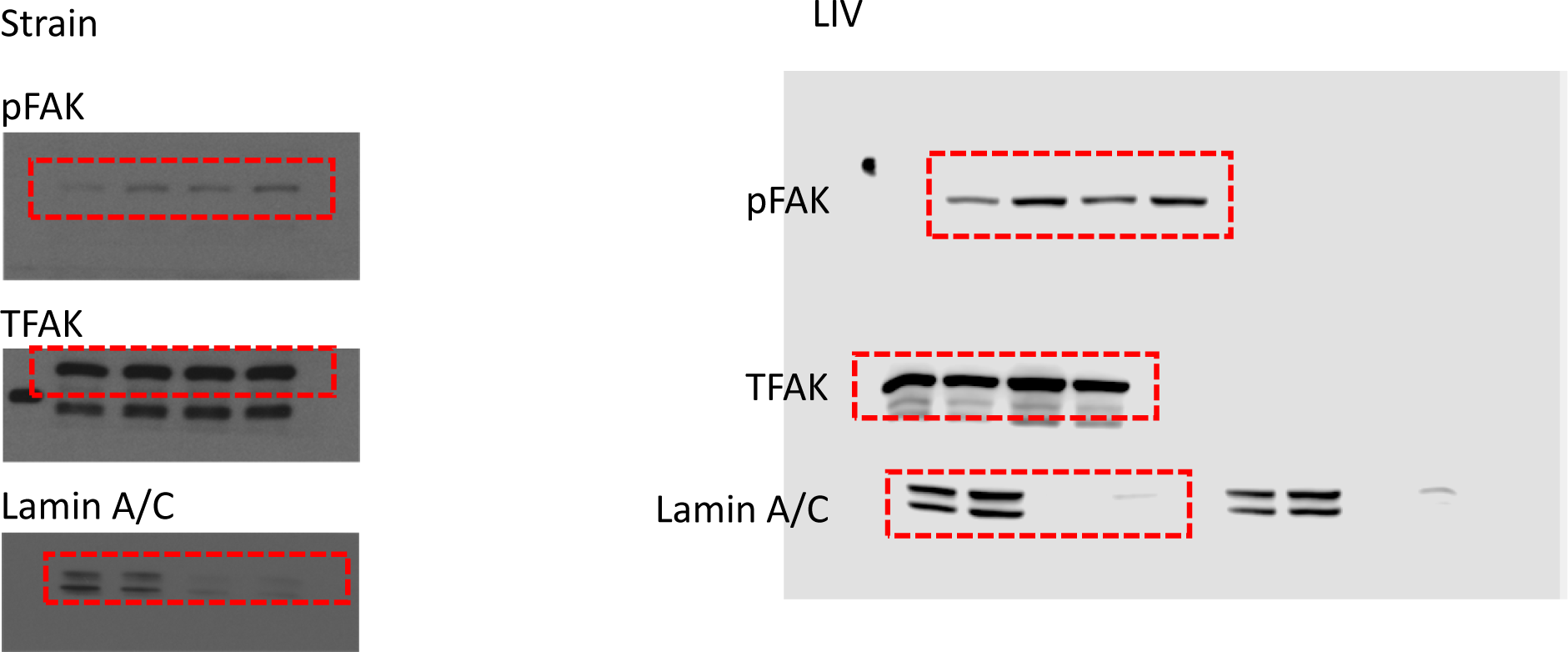
Unprocessed blots used in Figure 3 as obtained by LiCor C-DiGit blot scanner.

**Fig. S9.**
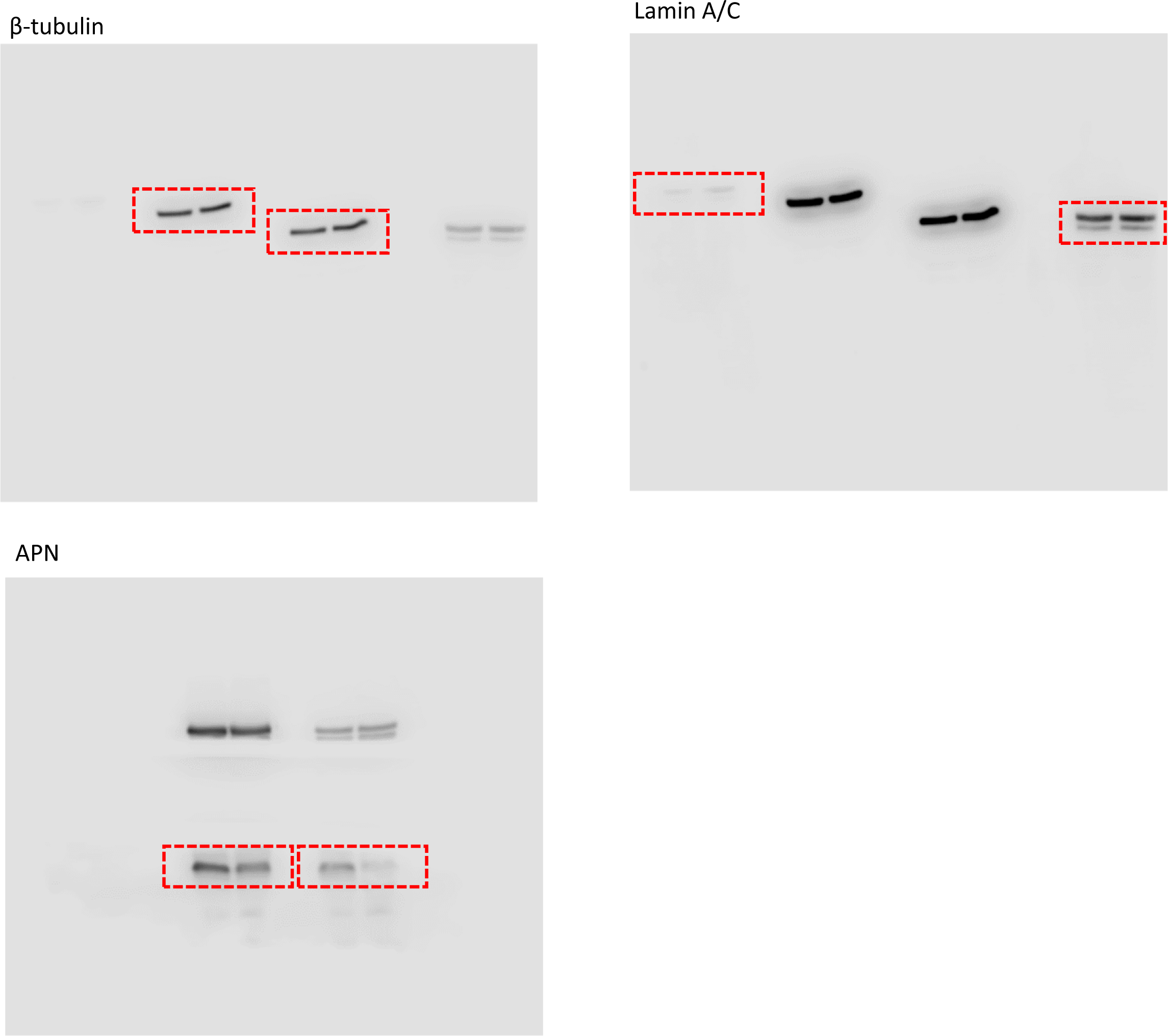
Unprocessed blots used in Figure 4 as obtained by LiCor C-DiGit blot scanner.

**Table S1:** Cell culture and pharmacological reagents and their final concentrations

| Cell culture and pharmacological reagents |  | Final Concentration |
| --- | --- | --- |
| IMDM | GIBCO | - |
| DMEM | Caisson Laboratories | - |
| FCS | Atlanta Biologicals | 10% v/v |
| Penicillin/streptomycin | GIBCO | 1% v/v |
| Dexamethasone |  | 0.1uM |
| Insulin |  | 5ug/ml |

**Table S2:**
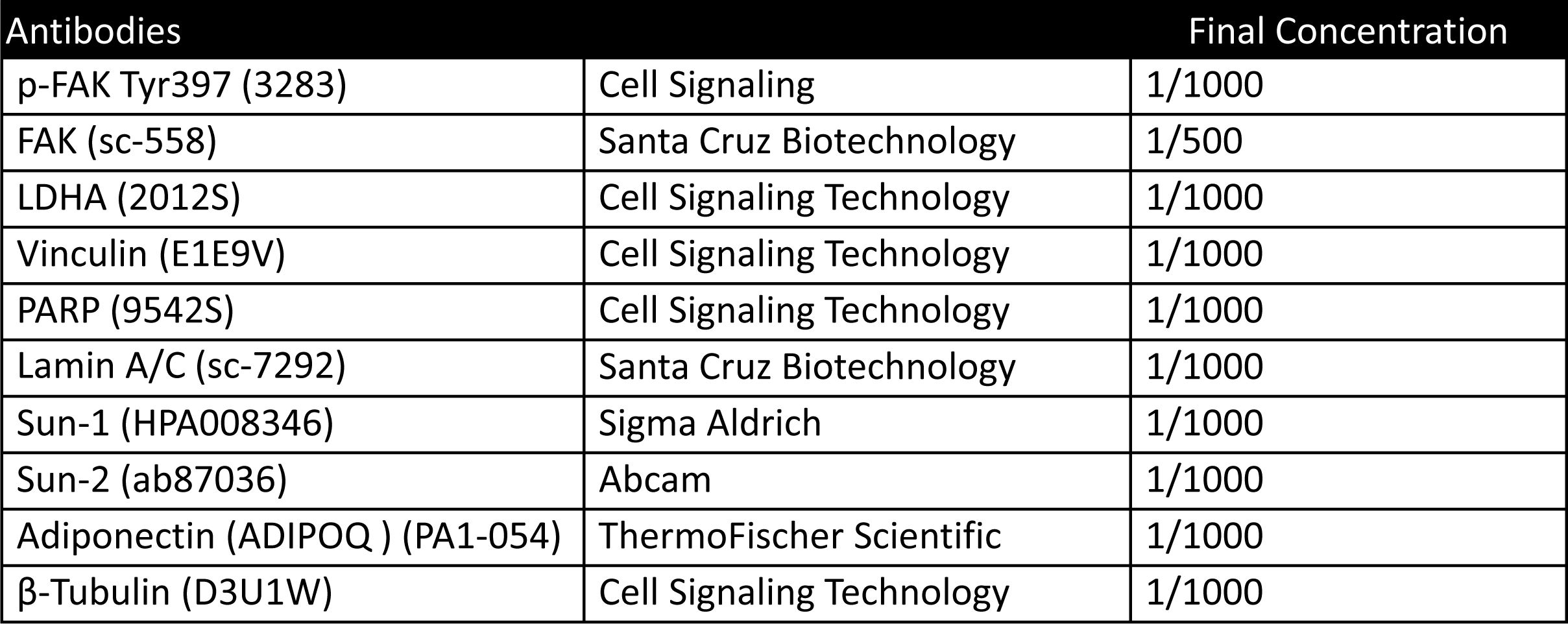
Antibodies used and their final concentrations for western blots.

**Table S3:** Immunostaining antibodies and reagents and their final concentrations.

| Immunostaining antibodies and Reagents |  | Final Concentration |
| --- | --- | --- |
| Hoechst 33342 | Thermo Scientific | 1 µg/mL |
| Alexa Fluor 488 Phalloidin | Life Technologies | 0.1µM |
| Lamin A/C (sc-7292) | Santa Cruz Biotechnology | 1:300 |
| Sun-1 (MABT892) | EMD Millipore | 1:300 |
| Sun-2 (IQ444) | Immuquest | 1:300 |

## References

1. Shumaker DK, Dechat T, Kohlmaier A et al. Mutant nuclear lamin A leads to progressive alterations of epigenetic control in premature aging. **Proceedings of the National Academy of Sciences**. 2006;103:8703–8708.

2. Schreiber KH, Kennedy BK. When lamins go bad: nuclear structure and disease. **Cell**.2013;152:1365–1375.

3. De Sandre-Giovannoli A, Bernard R, Cau P et al. Lamin a truncation in Hutchinson-Gilford progeria. **Science (New York, NY**). 2003;300:2055.

4. Constantinescu D, Gray HL, Sammak PJ et al. Lamin A/C expression is a marker of mouse and human embryonic stem cell differentiation. **STEM CELLS**. 2006;24:474–474.

5. Rober RA, Weber K, Osborn M. Differential timing of nuclear lamin A/C expression in the various organs of the mouse embryo and the young animal: a developmental study. **Development (Cambridge**, **England**). 1989;105:365–378.

6. Stewart C, Burke B. Teratocarcinoma stem cells and early mouse embryos contain only a single major lamin polypeptide closely resembling lamin B. **Cell**. 1987;51:383–392.

7. Underwood JM, Becker KA, Stein GS et al. The Ultrastructural Signature of Human Embryonic Stem Cells. **J Cell Biochem**. 2017;118:764–774.

8. Lammerding J, Fong LG, Ji JY et al. Lamins A and C but not lamin B1 regulate nuclear mechanics. **The Journal of biological chemistry**. 2006;281:25768–25780.

9. Lee JS, Hale CM, Panorchan P et al. Nuclear lamin A/C deficiency induces defects in cell mechanics, polarization, and migration. **Biophysical journal**. 2007;93:2542–2552.

10. Lammerding J, Schulze PC, Takahashi T et al. Lamin A/C deficiency causes defective nuclear mechanics and mechanotransduction. **The Journal of Clinical Investigation**. 2004;113:370–378.

11. Verstraeten VL, Ji JY, Cummings KS et al. Increased mechanosensitivity and nuclear stiffness in Hutchinson-Gilford progeria cells: effects of farnesyltransferase inhibitors. **Aging cell**. 2008;7:383–393.

12. Sullivan T, Escalante-Alcalde D, Bhatt H et al. Loss of A-type lamin expression compromises nuclear envelope integrity leading to muscular dystrophy. **J Cell Biol**. 1999;147:913–920.

13. Sabatelli P, Lattanzi G, Ognibene A et al. Nuclear alterations in autosomal-dominant Emery-Dreifuss muscular dystrophy. **Muscle & Nerve**. 2001;24:826–829.

14. Bronshtein I, Kepten E, Kanter I et al. Loss of lamin A function increases chromatin dynamics in the nuclear interior. **Nature Communications**. 2015;6:8044.

15. Ranade D, Pradhan R, Jayakrishnan M et al. Lamin A/C and Emerin depletion impacts chromatin organization and dynamics in the interphase nucleus. **BMC Molecular and Cell Biology**. 2019;20:11.

16. Stephens AD, Banigan EJ, Marko JF. Separate roles for chromatin and lamins in nuclear mechanics. **Nucleus (Austin**, **Tex**). 2018;9:119–124.

17. Moghadam FH, Tayebi T, Dehghan M et al. Differentiation of bone marrow mesenchymal stem cells into chondrocytes after short term culture in alkaline medium. International journal of hematology-oncology and stem cell research. 2014;8:12–19.

18. Sen B, Xie Z, Case N et al. Mechanical signal influence on mesenchymal stem cell fate is enhanced by incorporation of refractory periods into the loading regimen. **Journal of biomechanics**. 2011;44:593–599.

19. Sen B, Xie Z, Case N et al. Mechanical strain inhibits adipogenesis in mesenchymal stem cells by stimulating a durable beta-catenin signal. **Endocrinology**. 2008;149:6065–6075.

20. Sen B, Guilluy C, Xie Z et al. Mechanically induced focal adhesion assembly amplifies anti-adipogenic pathways in mesenchymal stem cells. **Stem Cells**. 2011;29:1829–1836.

21. Baskan O, Mese G, Ozcivici E. Low-intensity vibrations normalize adipogenesis-induced morphological and molecular changes of adult mesenchymal stem cells. **Proceedings of the Institution of Mechanical Engineers Part H**, **Journal of engineering in medicine**. 2017:954411916687338.

22. Uzer G, Pongkitwitoon S, Ete Chan M et al. Vibration induced osteogenic commitment of mesenchymal stem cells is enhanced by cytoskeletal remodeling but not fluid shear. **Journal of biomechanics**. 2013;46:2296–2302.

23. Bas G, Loisate S, Hudon SF et al. Low Intensity Vibrations Augment Mesenchymal Stem Cell Proliferation and Differentiation Capacity during in vitro Expansion. **Scientific reports**. 2020;10:9369.

24. Karadas O, Mese G, Ozcivici E. Low magnitude high frequency vibrations expedite the osteogenesis of bone marrow stem cells on paper based 3D scaffolds. **Biomedical Engineering Letters**. 2020;10:431–441.

25. Akter R, Rivas D, Geneau G et al. Effect of lamin A/C knockdown on osteoblast differentiation and function. **Journal of bone and mineral research : the official journal of the American Society for Bone and Mineral Research**. 2009;24:283–293.

26. Oldenburg A, Briand N, Sørensen AL et al. A lipodystrophy-causing lamin A mutant alters conformation and epigenetic regulation of the anti-adipogenic MIR335 locus. **The Journal of cell biology**. 2017;216:2731–2743.

27. Bermeo S, Vidal C, Zhou H et al. Lamin A/C Acts as an Essential Factor in Mesenchymal Stem Cell Differentiation Through the Regulation of the Dynamics of the Wnt/β-Catenin Pathway. **Journal of Cellular Biochemistry**. 2015;116:2344–2353.

28. Li W, Yeo LS, Vidal C et al. Decreased bone formation and osteopenia in lamin a/c- deficient mice. **PloS one**. 2011;6:e19313.

29. Geiger B, Spatz JP, Bershadsky AD. Environmental sensing through focal adhesions. **Nature Reviews Molecular Cell Biology**. 2009;10:21–33.

30. Hamadi A, Bouali M, Dontenwill M et al. Regulation of focal adhesion dynamics and disassembly by phosphorylation of FAK at tyrosine 397. **Journal of Cell Science**. 2005;118:4415–4425.

31. Uzer G, Thompson WR, Sen B et al. Cell Mechanosensitivity to Extremely Low- Magnitude Signals Is Enabled by a LINCed Nucleus. **STEM CELLS**. 2015;33:2063–2076.

32. Uzer G, Fuchs RK, Rubin J et al. Concise Review: Plasma and Nuclear Membranes Convey Mechanical Information to Regulate Mesenchymal Stem Cell Lineage. **Stem Cells**. 2016;34:1455–1463.

33. Sen B, Xie Z, Case N et al. mTORC2 regulates mechanically induced cytoskeletal reorganization and lineage selection in marrow-derived mesenchymal stem cells. **Journal of bone and mineral research : the official journal of the American Society for Bone and Mineral Research**. 2014;29:78–89.

34. Thompson WR, Yen SS, Uzer G et al. LARG GEF and ARHGAP18 orchestrate RhoA activity to control mesenchymal stem cell lineage. Bone. 2018;107:172–180.

35. Tontonoz P, Spiegelman BM. Fat and Beyond: The Diverse Biology of PPARγ. **Annual Review of Biochemistry**. 2008;77:289–312.

36. Pagnotti GM, Styner M, Uzer G et al. Combating osteoporosis and obesity with exercise: leveraging cell mechanosensitivity. **Nature reviews Endocrinology**. 2019;15:339–355.

37. Uzer G, Bas G, Sen B et al. Sun-mediated mechanical LINC between nucleus and cytoskeleton regulates βcatenin nuclear access. **Journal of biomechanics**. 2018;74:32–40.

38. Sen B, Xie Z, Case N et al. Mechanical signal influence on mesenchymal stem cell fate is enhanced by incorporation of refractory periods into the loading regimen. **Journal of biomechanics**. 2011;44:593–599.

39. Metsalu T, Vilo J. ClustVis: a web tool for visualizing clustering of multivariate data using Principal Component Analysis and heatmap. **Nucleic acids research**. 2015;43:W566–570.

40. Lee K, Um SH, Rhee DK et al. Interferon-alpha inhibits adipogenesis via regulation of JAK/STAT1 signaling. **Biochimica et biophysica acta**. 2016;1860:2416–2427.

41. Peister A, Mellad JA, Larson BL et al. Adult stem cells from bone marrow (MSCs) isolated from different strains of inbred mice vary in surface epitopes, rates of proliferation, and differentiation potential. **Blood**. 2004;103:1662–1668.

42. Dobin A, Davis CA, Schlesinger F et al. STAR: ultrafast universal RNA-seq aligner. **Bioinformatics (Oxford**, **England**). 2013;29:15–21.

43. Love MI, Huber W, Anders S. Moderated estimation of fold change and dispersion for RNA-seq data with DESeq2. **Genome biology**. 2014;15:550.

44. Dudakovic A, Camilleri E, Riester SM et al. High-resolution molecular validation of self- renewal and spontaneous differentiation in clinical-grade adipose-tissue derived human mesenchymal stem cells. **J Cell Biochem**. 2014;115:1816–1828.

45. Wang H, Rodríguez A. Identifying pediatric cancer clusters in Florida using loglinear models and generalized lasso penalties. **Stat Public Policy (Phila**). 2014;1:86–96.

46. Dudakovic A, Camilleri ET, Paradise CR et al. Enhancer of zeste homolog 2 (Ezh2) controls bone formation and cell cycle progression during osteogenesis in mice. **The Journal of biological chemistry**. 2018;293:12894–12907.

47. Dennison E, Cooper C. Epidemiology of osteoporotic fractures. **Horm Res**. 2000;54 Suppl 1:58–63.

48. Kalari KR, Nair AA, Bhavsar JD et al. MAP-RSeq: Mayo Analysis Pipeline for RNA sequencing. **BMC bioinformatics**. 2014;15:224.

49. Kim D, Pertea G, Trapnell C et al. TopHat2: accurate alignment of transcriptomes in the presence of insertions, deletions and gene fusions. **Genome biology**. 2013;14:R36.

50. Anders S, Pyl PT, Huber W. HTSeq--a Python framework to work with high-throughput sequencing data. **Bioinformatics (Oxford**, **England**). 2015;31:166–169.

51. Raharjo WH, Enarson P, Sullivan T et al. Nuclear envelope defects associated with LMNA mutations cause dilated cardiomyopathy and Emery-Dreifuss muscular dystrophy. **J Cell Sci**. 2001;114:4447–4457.

52. Kim J-K, Louhghalam A, Lee G et al. Nuclear lamin A/C harnesses the perinuclear apical actin cables to protect nuclear morphology. **Nature communications**. 2017;8:2123.

53. Corne TDJ, Sieprath T, Vandenbussche J et al. Deregulation of focal adhesion formation and cytoskeletal tension due to loss of A-type lamins. **Cell Adh Migr**. 2017;11:447–463.

54. Chancellor TJ, Lee J, Thodeti CK et al. Actomyosin tension exerted on the nucleus through nesprin-1 connections influences endothelial cell adhesion, migration, and cyclic strain-induced reorientation. **Biophysical journal**. 2010;99:115–123.

55. Wang W, Shi Z, Jiao S et al. Structural insights into SUN-KASH complexes across the nuclear envelope. **Cell Research**. 2012;22:1440–1452.

56. Haque F, Lloyd DJ, Smallwood DT et al. SUN1 Interacts with Nuclear Lamin A and Cytoplasmic Nesprins To Provide a Physical Connection between the Nuclear Lamina and the Cytoskeleton. **Molecular and Cellular Biology**. 2006;26:3738–3751.

57. Liang Y, Chiu PH, Yip KY et al. Subcellular Localization of SUN2 Is Regulated by Lamin A and Rab5. **PloS one**. 2011;6:e20507.

58. Naito M, Omoteyama K, Mikami Y et al. Suppression of lamin A/C by short hairpin RNAs promotes adipocyte lineage commitment in mesenchymal progenitor cell line, ROB-C26. **Histochemistry and cell biology**. 2012;137:235–247.

59. Khan KA, Dô F, Marineau A et al. Fine-Tuning of the RIG-I-Like Receptor/Interferon Regulatory Factor 3-Dependent Antiviral Innate Immune Response by the Glycogen Synthase Kinase 3/β-Catenin Pathway. **Molecular and Cellular Biology**. 2015;35:3029–3043.

60. Smith JL, Jeng S, McWeeney SK et al. A MicroRNA Screen Identifies the Wnt Signaling Pathway as a Regulator of the Interferon Response during Flavivirus Infection. **Journal of Virology**. 2017;91:e02388–02316.

61. Cartwright T, Senussi O, Grady MD. The Mechanism of the Inactivation of Human Fibroblast Interferon by Mechanical Stress. Journal of General Virology. 1977;36:317-665 321.

